# Discovery of dynamic changes in 3D chromatin architecture through polymer physics model

**DOI:** 10.1101/2024.04.11.589000

**Authors:** Anubhooti, Wasim Abdul, Priyanka Kriti Narayan, Jagannath Mondal, Jagan Pongubala

**Affiliations:** University of Hyderabad; Tata Institute of Fundamental Research, Hyderabad; Tata Institute of Fundamental Research Hyderabad

**Keywords:** Chromatin organisation, gene expression, polymer modelling, 3D structure, coarse-graining, simulations

## Abstract

The 3D organisation of the genome provides an intricate relationship between the chromatin architecture and its effects on the functional state of the cell. Recent advances in high-throughput sequencing and chromosome conformation capture technologies elucidated a comprehensive view of chromatin interactions on a genome-wide scale but provides only a 2D representation of how the chromatin is organised inside the cell nucleus. To quantitatively understand the structural alterations and dynamics of chromatin in 3D, we have developed a computational model that not only captures the hierarchical structural organisation but also provides mechanistic insights into the dynamics of spatial rearrangements of chromatin in developing lymphoid lineage cells. From the combination of approaches of polymer physics representing chromatin as a homopolymeric chain and incorporation of the biological information of chromosomal interactions inferred from the Hi-C data, we generated a coarse grained bead-on-a-string polymer model of chromatin to comprehend the mechanisms underlying the differential chromatin architecture. Our study showed that our simulated chromatin structure recapitulates the intrinsic features of chromatin organisation, including the fractal globule nature, compartmentalization, presence of topologically associating domains (TADs), phase separation and spatial preferences of genomic regions in the chromosomal territories. Comparative analyses of these simulated chromatin structures of differentiating B cell stages revealed compartmental switching and changes in the spatial positioning of lineage specific genomic regions. Analysis of the compactness of the switched regions showed insights into their acquired open-closed states for gene regulation and hence governing the cell fate through consequent structural rearrangement. Based on the remarkable performance of our model, we emphasise on its predictive potential by identifying switching of novel regions that demonstrated undergoing structural rearrangement which was subsequently validated through their differential expression patterns *in vitro*. These results reveal that although the chromatin organisation seems similar in most cell types, it undergoes distinct structural changes for the regulatory role of chromatin in sustaining cell specificity.

## Introduction

Our genome is responsible for expressing genes in a regulated manner across hundreds of different cell types. This is quite evident from the fact that though all the cells of an organism have identical genetic blueprint, yet they are specialised to perform remarkably diverse functions across cell types. This is principally because of the differential expression & regulation of genomic regions in distinct cells through the cross-talk between the *cis*-regulome and various trans-regulatory elements such as the transcription factors (TFs). These interactions within distinct nuclear compartments are, in fact, mediated spatially via their three-dimensional organisation and positioning in nuclear space. The 3D structural arrangement of the genome, thus, facilitates the genomic interactions and plays a pivotal role in gene regulation, consequently governing the functional state and fate of the cell.

The 3D architecture of the genome varies with cell types and also the developmental stages (Liu *et al* 2018) which implies that the knowledge of chromosomal spatio-temporal arrangement and structural dynamics across cell types is the key to understanding its impact on differential gene expressions. Recent findings have also highlighted the importance of this structural order in controlling embryonic development and how its disruption may lead to human anomalies (Misteli 2010, Spielmann *et al* 2018, Lupiáñez *et al* 2016, Anania and Lupiáñez, 2020). Further, elaborative functional importance of 3D genome architecture is reviewed in Bickmore and Steensel 2013, Cavalli and Misteli 2013, Edelman and Fraser 2012. For these reasons, understanding this 3D organisation of the eukaryotic genome and exploring the fundamental question of how the spatial conformation of a chromosome affects genetic and biological functions, has become of paramount importance in biomedical research and life sciences in general.

The 3D organisation of the genome is a critical player in the intricate orchestration of cellular development, particularly during the transition from the pre-pro-B to pro-B cell stage. This transition involves a profound shift in cell identity, as the cell commits to a specific path of becoming a mature B lymphocyte. The genome’s spatial architecture plays a pivotal role in this process by ensuring that the right genes are activated or silenced at the right time and developmental stage. It enables the efficient communication between enhancer elements, which act like molecular switches, and the genes they regulate. This spatial conformation ensures that genes relevant to B cell development are brought into close proximity, while those unrelated are secluded, creating a finely tuned symphony of gene expression. Consequently, the 3D genome organisation orchestrates the complex developmental journey from the pre-pro-B to pro-B cell stage, allowing for precise regulation of gene expression and ultimately determining the cell’s fate in the immune system.

The emerging importance of chromatin organisation has inspired the development of a variety of new experimental techniques such as the imaging techniques (e.g. FISH) and biochemical techniques (e.g. use of combination of proximity ligation and high-throughput sequencing chromosome conformation capture (Lieberman-Aiden *et al* 2009) and related methods (Kalhor *et al* 2012)) for elucidating a comprehensive genome-wide view of chromatin interactions. These methods are able to provide information on the cellular interactome at a whole-genome scale. However, they only provide a static 2D representation of the genome and prevent a direct quantitative description of the folding, movement and interaction of chromosomes within the nucleus. The dynamics of architectural changes via the 3D spatial rearrangements of genomic regions during varied cellular conditions is yet to be explored in its entirety. The gaps are filled using computational methods that can reveal spatial relationships between genomic regions that are not directly visible in the underlying data. In order to apprehend the 2D genome-wide contact data and unravel the principles shaping chromosome 3D structure and its functional implications, models from polymer physics were employed (reviewed in Rosa and Zimmer 2014, Chiariello *et al* 2016). These models have proven to be an essential tool in dissecting the significant mechanisms underlying chromosome spatial organisation (Dekker and Mirny 2016). Based on the physical characteristics of chromatin, these models helped deduce the rules governing genome folding (Dekker et al 2013). For example, information on the folding state of individual chromosomes based on physical constraints driving polymer compaction (Barbieri *et at* 2013, Rosa and Everaers 2014, Naumova *et al* 2013), on the formation of chromosome territories from physical forces that prevent polymer mixing (Mateos-Langerak *et al* 2009, Bohn and Heermann 2011) and on chromosomal interactions (Cook and Marenduzzo 2009, Jerabek and Heermann 2012). Other models such as the strings and binders switch (SBS) model (Barbieri *et al* 2012), which was first put forth to describe a variety of basic behaviours of chromatin in living cells has recently been expanded to investigate hierarchical chromosome architectures (Fraser *et al* 2015) and the consequences of structural variations on chromatin architecture (Bianco *et al* 2018). However, homopolymer models that are solely guided by a limited number of physical assumptions and parameters with only geometrical and topological constraints (Grosberg *et al* 1988, Bohn *et al* 2007, Lieberman-Aiden *et al* 2009, Mirny 2011, Halverson *et al* 2014, Kang *et al* 2015) may not fully capture all of the physical principles behind chromosomal organisation. Although these *direct* models have helped in understanding qualitative & quantitative properties, they retained simplified physical assumptions failing to take into account every aspect of the extensive experimental data sets. In more recent years, transcriptional factor binding with chromosomes has been modelled where the chromatins have been modelled as block polymers with binding factors explaining how domains are created (Brackley *et al* 2016). Another method elucidated the formation of TADs and contact maps’ prediction upon deletion of CTCF-binding sites based on the understanding of the convergent orientation of the CTCF-binding motifs (Fudenberg *et al* 2016, Sanborn *et al* 2015). On the other hand, the *indirect* or *inverse* models depended on the rich experimental data sets, such as the genome-wide contact frequencies, as input in order to reconstruct the 3D structure of a genome through distance-fitting and multidimensional scaling (Zhang *et al* 2013, Lesne *et al* 2014). Such models lack in their predictive power, for example, it is impossible to foresee the effects of a translocation or a change in gene expression through these models since a new data set would be required from such experiments as an input for the reconstruction.

Another emerging approach is utilising data from Hi-C, FISH and epigenetic states in a copolymer-type model to construct a more accurate picture of 3D chromosomal structure (Jost *et al* 2014, Brackley *et al* 2016, Wang *et al* 2015, Szalaj *et al* 2016, Tjong *et al* 2016, Di Stefano *et al* 2016, Shi *et al* 2018). Limitations of both *direct* and *inverse* models led us to the development of our hybrid model, along the similar lines, where we generated a physical coarse-grained bead-on-a-string polymer model and incorporated the experimental datasets as an input and let it further evolve with time. We introduce here a computational model of a chromatin of differentiating B-cells on which we performed simulations after incorporation of the genome-wide chromatin interaction data of Hi-C in order to study cell type specific structural dynamics. From this time-evolved trajectories, we have studied the dynamic changes occurring in the two systems representing the two cell stages. We were able to derive a fundamental relationship between genome organisation & cell type-specific gene expression and regulation through the use of this combinatorial approach, which combined the concepts of polymer physics with minimal biological information significant enough for studying and providing mechanistic insights into cell-type specific 3D chromatin folding.

Through this work, we show the spatial dynamics in 3D in terms of transitional rearrangements in chromatin organisation, thereby, predicting consequential gene expression patterns upon cell differentiation. This was not possible through data-driven reconstruction-based modelling approaches which are limited to reconstruction of only static chromatin structures based on the input provided. We studied the organisation of a murine chromosome that shows crucial changes during B cell development. In order to study its dynamics, we have generated time-evolved conformations of a self-avoiding polymer chain with Hi-C interactions incorporated as weak harmonic bonds and performed Langevin dynamic simulations. Additionally, benefitting from our polymer-based predictive approach, we went ahead to make significant differential structural predictions through our simulated structures, which could not be captured in other high-throughput experiments but were detected by our simulated structures. Based on our results, we were able to demonstrate a crucial association between 3D genomic organisation and cell-type-specific gene expressions, which are substantially influenced by structural chromosomal dynamics.

## Methods

### Data Acquisition

The experimental data used in the present study was obtained from the in-house experiments of high-throughput Hi-C sequencing data published online (<u>GSE85858</u>) (Boya *et al* 2017). The Ebf1^-/-^ cells indicate the Pre-Pro-B stage while the Pro-B cell stage is represented by Rag2^-/-^ cells.

### Model Generation

We have computationally modelled the chromosome 11 of mouse genome as a beads-on-a-string homopolymeric chain consisting of spherical non-overlapping beads (self-avoiding walk polymer) of defined diameter (σ) connected by a spring. Each of the beads in the present model maps a genomic region of size 40kb, which is the same as the Hi-C matrix resolution as reported by Boya *et al* 2017. Considering the length of chromosome 11 in the mouse genome of 122082543bp (as in the mm10 genome build), the total number of beads in our model system turned out to be 3053 (122082543bp/40x10^3^bp). 200 such initial self-avoiding polymer chains with 3053 beads each, were generated where each polymer chain was defined in a confinement of radius (r_conf_) 0.986μm in real units. The confinement depicts the chromosomal territory (Figure 1), where the radius of the confinement is calculated proportional to the known genomic volume fraction of an eukaryotic cell (see SI). Based on the volume fraction of 0.1 for a eukaryotic genome (Chen *et al* 2017), the diameter (σ) of the 40kb sized spherical bead was calculated to be 63.13nm (see SI) and the radius of confinement to be 15.6 times larger than the bead diameter, i.e. r_conf_= 15.6σ (see SI). The contact information from the Hi-C data was then integrated to these initial random generated structures, the details of which are mentioned in the next section.

**Figure 1.**
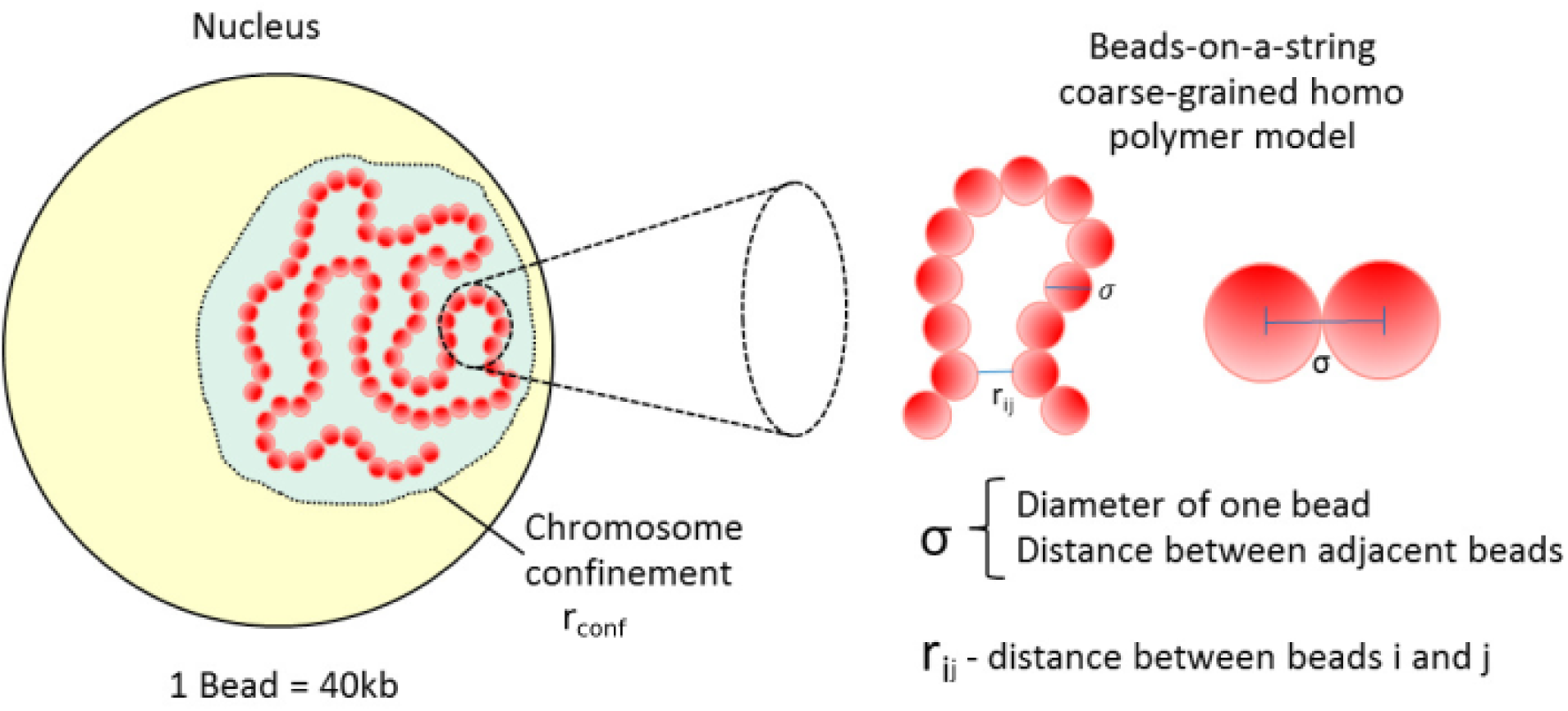
Prototype of initial configuration. Prototype representing the bead-on-a-string homopolymeric chromatin model with bead size (σ) and radius of confinement (r_conf_) shown. The size of one bead is considered to be 1σ or 40kb in genomic units.

The energetic interactions between these beads were modelled via bonded and nonbonded interactions, whose analytical forms have been detailed in SI. An excluded volume interaction is implemented via the repulsive part of Lennard Jones interaction between any two consecutive 40 kbp-sized beads modelled by harmonically restrained bonds while the same between three consecutive beads were modelled via an angular potential.

### Incorporation of the Hi-C Data

Once the polymer chains are generated with the model parameters (σ and r_conf_), the biological information from the Hi-C interactome data is incorporated, thus obtaining the current hybrid model. The intra-chromosomal interaction data for chromosome 11 was obtained from the in-house generated genome-wide Hi-C for Pre-Pro-B cells as well as the Pro-B cells (Boya *et al* 2017) and was integrated into all the 200 initial polymer models as weak harmonic bonds. For this, we first extracted the N×N intra-chromosomal matrices from the normalised genome-wide Hi-C contact frequency matrices for both the Pre-Pro-B and Pro-B cell stages. The steps to process the raw reads of the Hi-C data have already been discussed in Boya *et al* 2017 wherein the Iterative mapping module of hiclib (https://github.com/mirnylab/hiclib-legacy by Mirny lab) was used. After ICE (iterative correction and eigenvector decomposition) normalisation, the corrected contact frequency matrix was converted to contact probability matrix using the method previously employed in Zhang and Wolynes 2015, i.e.

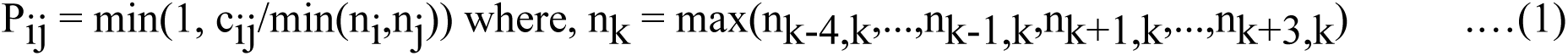

and c_ij_ is the contact frequency and P_ij_ is the contact probability between regions or beads

*i* and *j*. These experimentally derived Hi-C contact probability maps were integrated in the current model as harmonically restrained bonds between two given beads representing the corresponding 40kb sized genomic region in the Hi-C contact frequency matrix. However, unlike bonds between consecutive beads, these ‘Hi-C bonds’ between non-consecutive beads are restrained by contact probability-dependent distances and distance-dependent force constants. In particular, if the Hi-C pair interaction probability, P_ij_ between *i* & *j* of the N×N contact probability matrix is such that, P_ij_ >= P_c_ where P_c_ is the probability cut-off at 0.04, then the corresponding ‘Hi-C bond’ is modelled via a harmonic restraint of spring constant, (k_Hi-C_) defined as

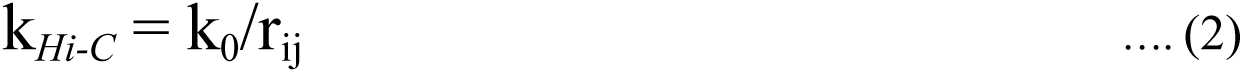

where, r_ij_ = σ/P_ij_, k_0_ = 2.0 kJ mol^-1^ nm^-2^. Here, the amplitude term k_0_, establishes the upper limit to the force constants of the Hi-C bonds. This function essentially implies weaker values of force constant for larger distances. The threshold probability cut-off P_c_, was chosen in order to consider only the minimal set of Hi-C data above the selected threshold of contact probability. An overview of the complete workflow is depicted in Figure 2.

**Figure 2.**
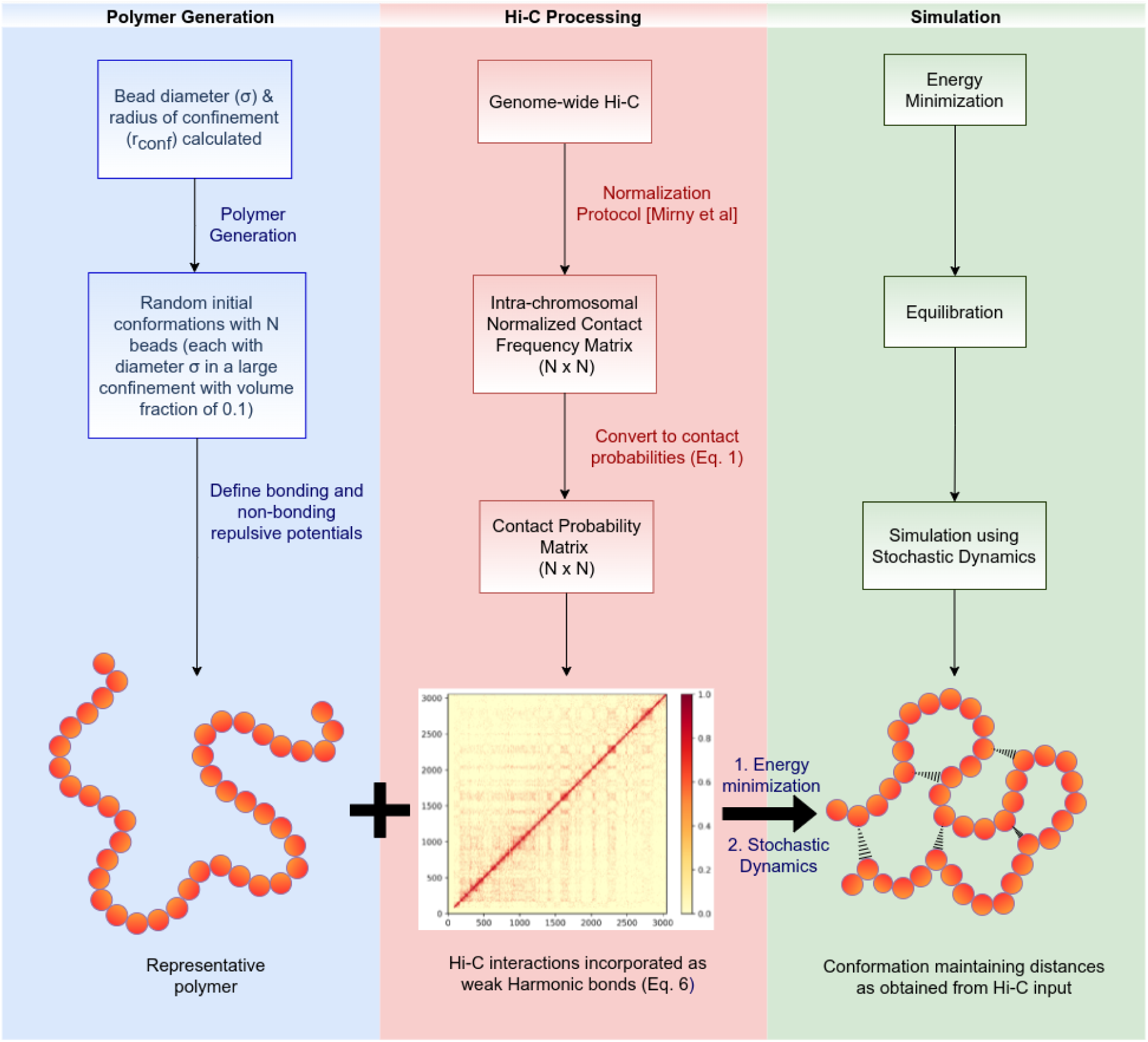
Schematic overview of the approach: The flow chart explains the workflow of the approach followed and the processes involved in each step.

### Simulation Details

All simulations are performed using the open source package GROMACS 5.0.7 (Abrahm *et al* 2015). We energy minimised the 200 polymer configurations, generated using the model described in the previous section, followed by 2×10^6^ steps of Langevin Dynamics simulation for each of these configurations. A Langevin thermostat set to 310K and a coupling constant of 1 ps was used to maintain the temperature of our system. Each of the 200 simulations were run for the total time of 2.372s in real units, 1 timestep(ts)=0.002𝛕 (see SI for derivation of time-scales) and at equal intervals of every 250 timesteps, we had saved the coordinates of the system. Thus, the number of configurations saved for each of the 200 simulations will be total timesteps/250, giving rise to 8000 simulation frames for each polymer configuration.

### Calculation of Simulated Contact Probability Matrix

For both Pre-Pro-B and Pro-B cell stages the simulations were done independently. Last 2000 frames of each of these 200 GROMACS trajectories were employed for any production analyses as these were energy minimised and had attained equilibrium. All inter-particle distances were calculated in order to generate a distance matrix for each of the 2000 frames. For a single trajectory, a final distance matrix (D) was then obtained by averaging over these 2000 total number of frames. We then generated a probability matrix (P) for each trajectory such that P = σ/D. Therefore, the probability matrix (P) is averaged over 2000 independent conformations in each trajectory. Further, a final simulation derived contact probability matrix (P_sim_) was generated by averaging over all the 200 trajectories. Therefore, the simulation-derived contact probability matrix (P_sim(PPB)_ for Pre-Pro-B and P_sim(PB)_ for Pro-B) is essentially averaged over 200×2000 independent conformations. Since a set of 2000 frames belong to a particular initial conformation, instead of generating 200 × 2000 = 4×10^5^ probability matrices and performing an average over them, we average over 200 matrices, which have already been averaged frame-wise. All the analysis was done by converting the distances to contact probabilities using MDAnalysis Python Package. We further filtered this simulated contact probability matrix by replacing elements in the matrix with zeros which were also zero in the experimental matrix to avoid any misinterpretations from our simulated matrix. To render the representative 3D conformations of the chromosome model, we have used the open-source package Visual Molecular Dynamics (VMD) (Humphrey *et al* 1996).

## Results

### Polymer-based model recapitulates chromosome-conformation capture data

To begin with, we have considered modelling chromosome 11 of mouse as this chromosome harbours crucial factors responsible for B cell development and also has a genomic length (122kb) which is intermediate in size. Hence, it is optimal in terms of handling complexity in a computationally affordable model. As described in the *Methods* section and in SI, we have modelled the chromosome as a beads-on-a-string homopolymer chain consisting of identical monomers or beads at 40kb resolution. The energy function incorporates experimentally rendered Hi-C probability matrix, excluded volume interaction (see Methods) where the resultant polymer model is constrained in a confinement that is commensurate with its chromosomal territory (Figure 1).

To validate our proposed computational model, we computed the simulation contact probability matrix by averaging over the ensemble of conformation simulated across multiple configurations (averaged over 200 × 2000 independent conformations) and compared it with the experimental contact matrix obtained from the Hi-C data (see SI for details of simulation contact probability calculation). Figure 3a compares the heatmap between simulation derived contact probability matrix and the experimental Hi-C contact probability matrix of chromosome 11 for Pre-Pro-B cell. We represent the same for Pro-B cell in SI (Figure S1a). With a Pearson correlation coefficient of 0.91 and 0.92 between corresponding experimental and simulated contact probability matrices of Pre-Pro-B and Pro-B cells, respectively, our model clearly indicates a very good agreement between simulations and experimental data for both cell stages. This is also evident through the remarkably similar checker-board patterns of the corresponding matrices in both cell types. The results also show that in spite of considering only a small percentage of experimental interactions above a threshold, P_c_, such that P_ij_ ≥ P_c_ where Pc = 0.04 in our simulations, our model faithfully reproduces not only the considered Hi-C interactions but also those experimental Hi-C interactions which weren’t included in the initial incorporation while generating the model. This result contributes to the model’s efficiency and performance with only limited input information. In the heatmap generated in Figure 3a, we also observed intense diagonal regions which indicate smaller distances having a higher contact probability between neighbouring chromosomal regions. This is justified since the proximal regions represented by the diagonal, tend to exhibit higher contact probabilities than the distal genomic regions that exhibit relatively smaller contact probabilities unless there is a possibility of formation of highly interacting regions such as TADs.

**Figure 3:**
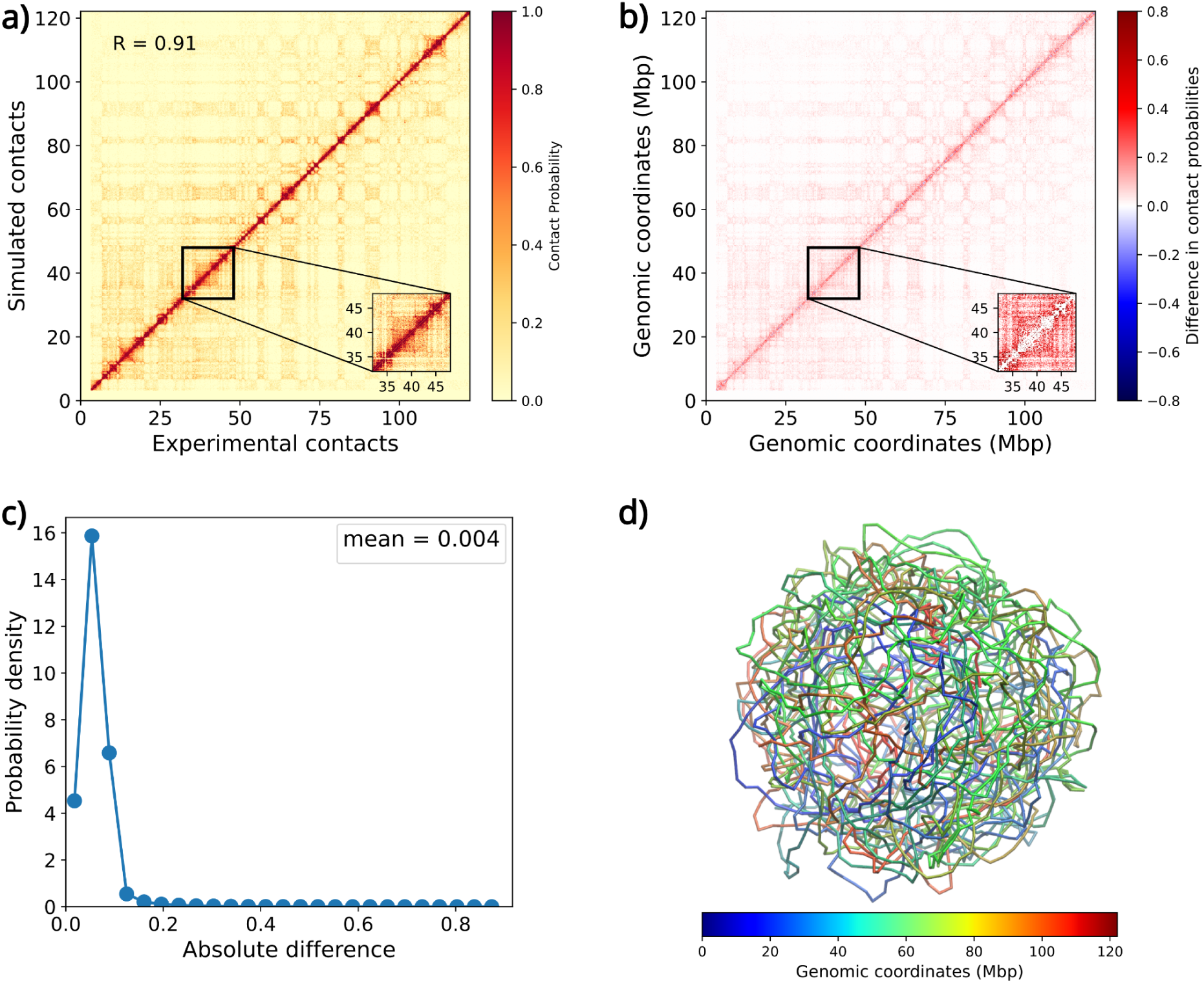
a) Comparison of experimental versus simulation-derived probabilities: The heatmap shows the comparison between experimental and simulated contact probabilities maps of chromosome 11 at 40kb resolution. b) Difference plot: The heatmap shows the difference between experimental and simulated contact probabilities maps of chromosome 11 at 40kb resolution. The blue and red colours in the colour bar indicate higher contact probability in the experimental data and simulations respectively. **c)** Absolute difference plot: The absolute difference plot shows maximum difference value of 0.004 between experimental and simulation-derived probabilities. **d)** Representative snapshot of a Pre-Pro-B chromosome. The regions of the chromosome have been coloured with respect to their genomic location.

Further, for a more rigorous assessment of computed simulation-derived matrix, we plotted a heatmap of the difference matrix calculated as difference between the simulation-derived contact probability matrices and the experimental contact probability matrices for both Pre-Pro-B (Figure 3b) and Pro-B (Figure S1b) cell stage. We observe that this difference for any bead *i* and *j* is minimal in both cell types, (the white regions in the difference plots), except in regions near the diagonal. In the diagonal region, simulation contact probability is estimated to be higher than the corresponding experimental probabilities. This could arise due to the high interaction frequencies between consecutive beads owing to their physical proximity that accounts for the over estimation of contact probabilities. Even in the absence of any contact experimental information, the simulation contact probabilities tend to be greater due to the closely packed adjacent beads as a result of the imposed confinement that leads to the observed difference. Thus, the results from the difference heatmap indicate that for longer genomic distances, the simulation-derived contact probabilities are in agreement and exhibit least difference with the experimental contact probabilities.

In order to quantify this difference, we generated the probability density plot of absolute differences between the experimental and simulation contact probabilities for both cell types (Figure 3c for Pre-Pro-B and Figure S1c for Pro-B) after excluding the noise near the diagonal regions that was due to the higher differences in the contact probabilities, as observed in Figure 3b and S1b. In the distribution of absolute values of the difference in contact probabilities between experiment and simulation data obtained from the difference heatmap, we see that the discrepancy between the simulation and experiment contact probability is as small as <0.1, indicating that there is a fair amount of agreement between the simulation and experiment contact probabilities. We observe that above 90% of the contact probabilities show the absolute difference close to 0, indicating almost negligible difference between the experimental and simulation probabilities, thereby, suggesting our model’s conformity with experiments. We observe the maximum difference value to be as low as 0.004 and 0.006 between experimental and simulation-derived probabilities for Pre-Pro-B (Figure 3c) & Pro-B cells (Figure S1c), respectively. Due to these small scale differences, we demonstrate that our model is sufficiently robust to be explored for investigating and predicting key chromosomal properties.

Finally, we also show a conformation representative from the ensemble of conformations for the structure of chromosome 11 obtained via simulations for Pre-Pro-B (Figure 3d) and Pro-B (Figure S1d). The snapshots were generated using VMD software and rendered for image quality purposes. The chromosomal regions have been coloured with respect to their genomic location for both Pre-Pro-B and Pro-B chromosome models.

### Model independently demonstrates chromatin hierarchical organisation

The previous results were obtained based on the initial input of the biological data provided during the model generation and incorporation of the Hi-C step as mentioned in *Methods*. In order to test the predictability, reliability and behaviour of our simulated model structures, we extended our investigation to explore some of the intrinsic properties of the chromatin which were independent of any implicit or explicit experimental inputs other than the Hi-C data used during the generation of the model. Beginning with the highest chromosomal level of organisation, we first investigated the nature of folding of our simulated chromatin structures. The standard method is by observing the scaling of contact probabilities P(*s*) as a function of genomic distance (*s*) which follows a power-law relationship also represented from its slope (Lieberman-Aiden *et al* 2009). To examine this relationship in our model, we plotted the intra-chromosomal contact probabilities as a function of genomic distance for both the simulated structures that represented the respective cell types (Figure 4a and S2a). The inverse power law scaling with the slope of -0.86 (*s^-0.86^)* and -0.83 (*s^-0.83^)* was observed in case of Pre-Pro-B (Figure 4a) and Pro-B cells (Figure S2a), respectively. This is in agreement with the already reported value of -1 (s^-1^) in case of fractal nature of the chromosome (Lieberman-Aiden *et al* 2009) thereby indicating that the folding and local packing of our simulated polymer, representing a single chromosome, also shows a fractal behaviour.

**Figure 4:**
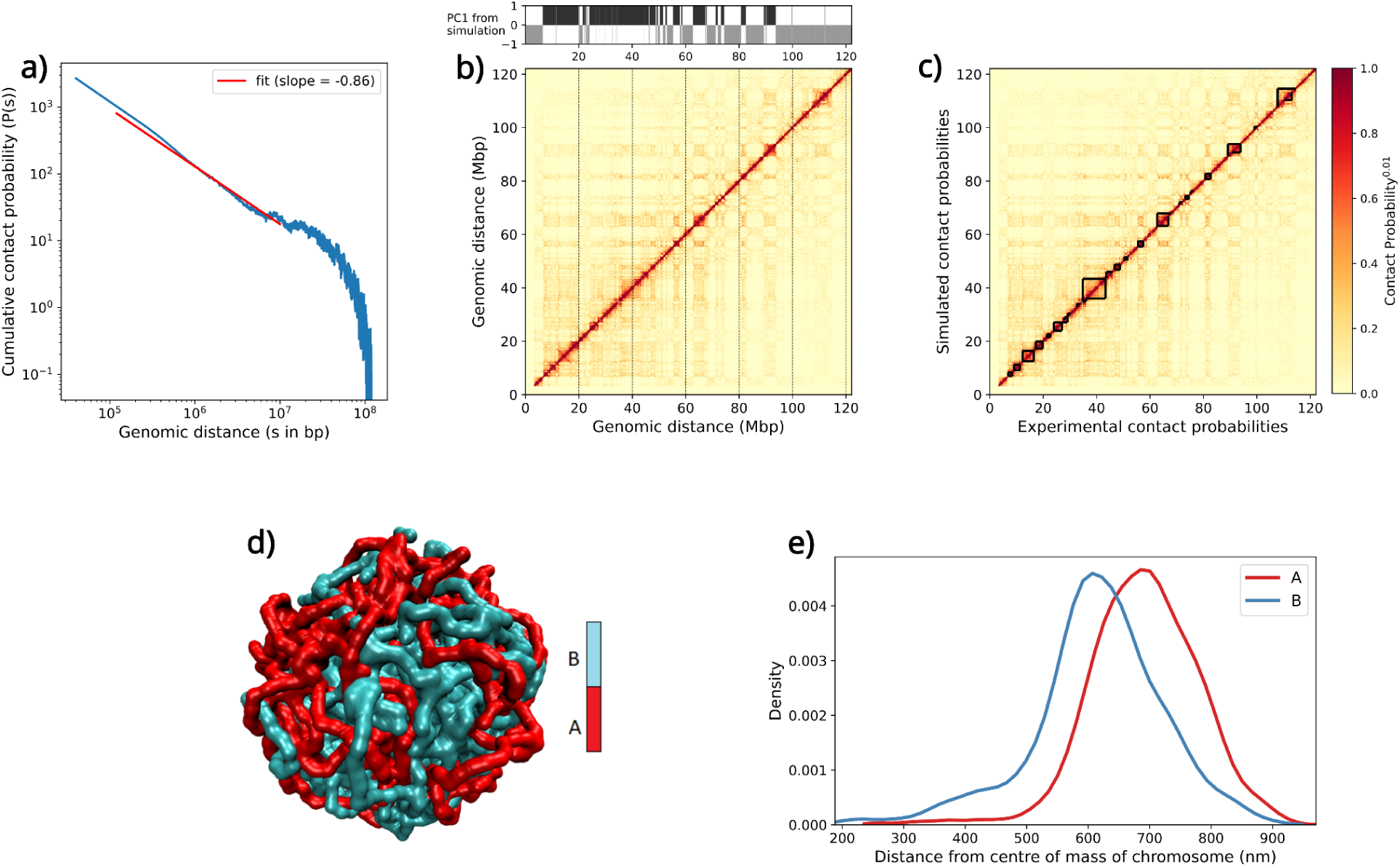
a) Chromatin folding prediction. Plot of contact probability as a function of genomic distance with a slope (fit shown in red) of -0.86 for Pre-Pro-B which is close to the slope of -1.0 for a fractal globule structure. **b) Prediction of chromatin state** Prediction of chromatin states from PCA (PC1 values) of the simulation derived contact probabilities is compared with the heatmap of experimental contact probabilities for Pre-Pro-B. The plot shows that the prediction of A compartments (or permissive regions) from the positive PC1 values (black region) in the simulation corresponds to the high contact probabilities in the experimental matrix while the prediction of B compartments (or repressive regions) from the negative PC1 values (grey region) in the simulation corresponds to the low contact probabilities in the experimental matrix. **c) Prediction of Topologically Associated Domains** We have used Armatus (Filippova *et al* 2015) to predict TADs from the simulated (blue triangles) structures and the experimental (green triangles) contacts obtained from Hi-C. The results are shown for Pre-Pro-B simulated structures. In this figure TADs which are at least 1 Mbp long have been shown. **d) Phase separation** Phase separation of A and B compartments in the simulated structures for Pre-Pro-B cell stage. Similar compartment regions (A compartment in red and B compartment in cyan colour) co-localize leading to a phase separation of active (permissive) from inactive (repressive) regions. **e) Preferential spatial positioning of compartments** Mean distances of A and B compartments from the centre of mass (COM) of the simulated structures of Pre-Pro-B. A compartments have larger mean distance from the COM of the chromatin indicating their preferential positioning farther from the centre, at the chromosomal periphery while smaller mean distances of B compartments indicates that their preferential positioning is in the interior of the chromatin, closer to the centre of the chromatin polymer.

Further at the next hierarchical chromatin organisation level, we investigated if our model can determine chromatin states into transcriptionally active and inactive regions, i.e. A and B compartments, essentially corresponding to the euchromatin and heterochromatin regions respectively. It has been shown previously that Principal Component analysis (PCA) is the canonical and the most popular method for identifying compartmental status of a given region where the first principal component (PC1) or eigenvector capturing the dimension with the highest variance, is used to to assess the region’s A/B compartmental status. The PC1 has been divided into two sets of values that correlates well with the corresponding regions in the genome as transcriptionally active or permissive and inactive or repressive regions wherein the positive PC1 values represent the permissive A compartment regions and negative values represent the repressive B compartment regions. (Lieberman-Aiden *et al* 2009). We implemented the same concept of PCA to our simulation results and overlaid it with the experimental contact probability heatmap as shown in Figure 4b and S2b. The results clearly demonstrate the segregation of the chromosome into A (active or euchromatin) and B (inactive or heterochromatin) compartments where the positive PC1 values of the simulation data correlate to the A compartment in the experimental heatmap while the negative PC1 values correspond to the B compartment in the experimental heatmap in case of both Pre-Pro-B (Figure 4b) and Pro-B (Figure S2b) cell types. This invariably confirms the correctness of our model wherein the chromatin status of different regions in the simulated chromatin model is predicted correctly in accordance with the corresponding chromatin status of those regions observed experimentally.

After the encouraging performance of our model at the sub-chromosomal level, we were interested to examine its behaviour at the sub-megabase level also. At this level, the chromatin is compacted and organised into highly self-interacting regions called Topologically Associated Domains (TADs) where TAD boundaries are important in gene regulation. TAD boundaries are crucial in insulating and preventing neighbouring regions from interacting with each other while orchestrating proper enhancer-promoter activity and cell-specific gene expression within TADs. These can be seen as ‘triangles’ near the diagonal in the Hi-C contact heatmap. Owing to such a critical role in gene regulation, we were keen if our model could represent TADs and TAD boundaries accurately. There are a number of well-established TAD prediction tools, such as Arrowhead (Rao et al 2014), TADbit (Serra et al 2017), TADtree (Weinreb and Raphael 2016), TopDom (Shin et al 2016) and many others. Based on the evaluation of many TAD callers and eventually choosing the one that produced the most consistent and visually pronounced TADs, we decided to use Armatus TAD caller (Filippova *et al* 2014) for our analysis.

In Armatus, TADs are defined using algorithms that detect switches in the directionality of interactions. Figure 4c shows the results for TAD calling for Pre-Pro-B and Figure S2c for Pro-B cells that compares the results of the simulated structures with their corresponding experimental data. We observe that the number of domains predicted for simulated structure is 480 and 463 for experimental data in case of Pre-Pro-B while for Pro-B, the number of domains predicted for simulated structure is 511 and 410 for the experimental data. The positions of the corresponding TADs in simulation versus experimental data for both the cell types is remarkably similar as shown in Figure 4c and Figure S2c. This is a *bonafide* agreement of the simulated chromatin structure even at such small sub-megabase pair level.

### Phase separation

Further, we examined for possible phase separation of A and B compartments in 3D space in our simulated structures. The dynamic phase separation of the genome had been proposed due to the flexible chromatin structure and movements (Hnisz *et al* 2017). Due to the respective spatial constraints to allow for differences in interactions of transcriptionally active and inactive regions with other genomic regions, these similar-state chromatin regions tend to co-localize and become phase separated. We tried to investigate the phase separation in Figure 4d for Pre-Pro-B and Figure S2d for Pro-B simulated structures and show that the similar compartment regions (A compartment in red and B compartment in cyan colour) co-localize leading to a phase separation of active (permissive) from inactive (repressive) regions. We generated these images by visualising the simulated structures in VMD. Although these results are after qualitative visual inspection only, it will be further interesting to observe differential patterns in these phase-separated compartments which we speculate to largely determine dynamic genome organisation and contribute towards cell fate decisions.

### Spatial positioning

To further quantitatively assess the preferential spatial positioning of these phase separated compartments in 3D space, we computed the mean distances of A and B compartments from the centre of mass (COM) of the simulated structures. The resultant plot in Figure 4e for Pre-Pro-B simulated structure and Figure S2e for Pro-B clearly shows that the A compartments have larger mean distance from the COM of the chromatin polymer indicating that these active regions tend to position themselves farther from the centre and towards the periphery of the chromatin. This positioning in 3D space in the exterior surface of the chromatin would allow easy accessibility of the genes harboured by these compartments to the transcriptional machinery of the cell, for their expression. On the other hand, the B compartments have smaller mean distance from the COM of the chromatin polymer indicating that these inactive regions are closer to the COM and are buried in the interior of the chromatin correlating to the inactivation of genes in those compartments. We can, thus, say that the phase separated compartments have preferential positioning in space which is directly related to their gene expression. Therefore, it can be deduced that our model’s predictions on the state and positioning of its chromatin regions in 3D space are in-line with the theoretical phenomenon where the spatial arrangement of chromatin has a significant influence on the genome function.

Taken together, our findings show that our model is able to successfully capture and predict some of the very important characteristic features of chromatin architecture at different levels of chromatin organisation, such as folding of chromatin as a fractal globule, transcriptional state of chromatin resulting into compartmentalization into A/B compartments, formation of TADs and prediction of TAD boundaries, phase separation of similar chromatin state regions and the spatial positioning of the transcriptionally variable regions in context of the chromatin, all of which are independent of any biological inputs other than a small subset of the Hi-C interactions used during model generation and are entirely the resultant properties and behaviour of our generated simulated structures. Hence, we have established the reliability and predictability of our model. We, now, use these characteristics as our model’s strengths and extend our investigation to further carry out comparative analysis of the two cell types and examine cell type specific differential changes.

### 3D modularity of chromatin

We have also examined the effect of deletion of regions and compared those partial chromatin regions with the entire chromosome in order to better understand the spatial modularity in chromatin folding. To do so, we considered different sizes of the chromatin polymer chains of N = 51, 101, 201, 501, 1001 and 2001 beads and simulated these self-avoiding chromatin chains of different lengths. These regions of variable lengths were selected starting from the centre of the chromatin polymer and extending at equal intervals towards the left and right of the chain. We estimated the radius of gyration, R_g_, for these various values of N in order to derive the power law scaling from our simulations. The resulting plot is shown in Figure 5a. By taking into account how, for large values of N, the polymer’s radius of gyration R_g_ behaves, one may apply the most popular critical exponent, the compactness index, ν, which has been studied previously for the three polymeric phases (Chernodub *et al* 2011). The value of ν corresponds to 1/2 for a random-walk polymer, 3/5 for self-avoiding walk polymers without any restraints and 1/3 if the self-avoiding chain polymer is in a collapsed phase inside a confined boundary. From our results in Figure 5a, we find that the chromatin behaves in a modular fashion with its R_g_ correlated as the slope corresponding to 0.297 which is slightly less than 1/3 for the collapsed state of self-avoiding walk polymer in a confinement. This is due to the presence of intra-chromosomal local interactions obtained from Hi-C that have been incorporated as weak harmonic bonds. Hence, the value (of 0.297) is little less than the expected value of 0.33. It is deduced that even though our chromatin polymer faithfully follows the polymeric properties, it is influenced slightly due to the presence of genomic interactions that in turn govern its overall folding. This essentially implies chromatin folding in a bad solvent having more intra-polymeric interactions than polymer-solvent interactions which is certainly the case since the chromatin-chromatin interactions are more prevalent in deciding the chromatin organisation and arrangement inside the nucleus. Since, we see that these interactions play a crucial role in governing the folding and dynamics of chromatin polymer, we were interested to investigate their nature of impact. For this, we plotted the R_g_ of different regions of the same size (=200 beads) and examined its behaviour. From Figure 5b, we show that short-ranged local interactions are more prevalent than the long-ranged interactions. Also, there is heterogeneity in these interactions as is evident from the different values of R_g_ for the same size of polymer. It is the presence of these heterogeneous local interactions that impacts the chromatin folding and three dimensional architecture of chromatin.

**Figure 5:**
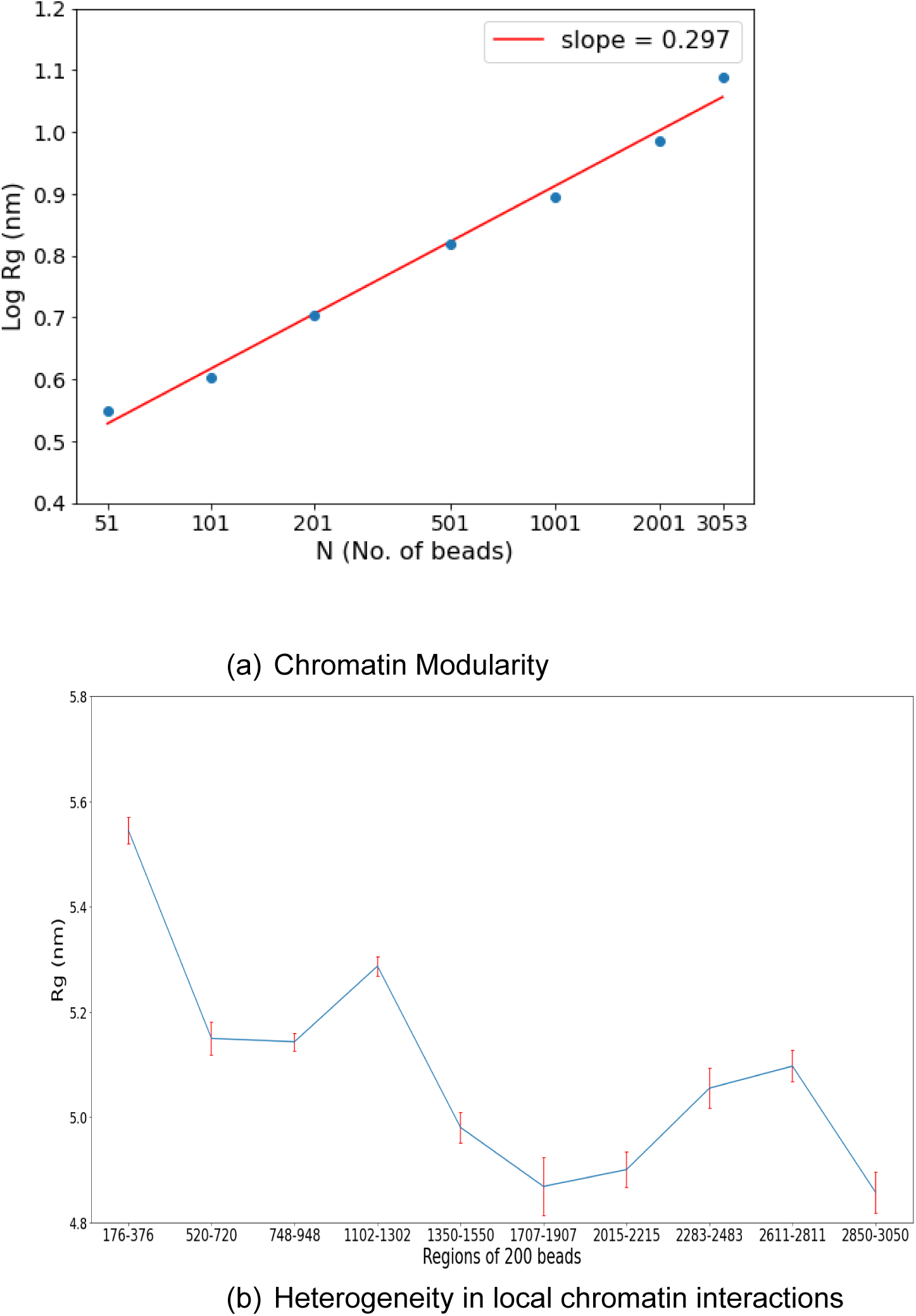
3D Chromatin Modularity. (a) Behaviour of chromatin regions in terms of their R_g_ values for different sizes of the polymeric chain. (b) Behaviour in terms of mean R_g_ values of different chromatin regions of same size. Standard deviation for each region is indicated as red error bars.

### Comparative analysis of simulated structures showed cell-type specific compartment switching

The remarkable agreement of the chromatin interactions, folding behaviour, compartmentalization and TAD formations between the simulated structures and the experimental data of the two cell stages, Pre-Pro-B and Pro-B, led us to extend the model’s usage in comparing the chromatin organisation and capturing the structural alterations during cell differentiation that could not be captured in experiments. We proceeded to specifically probe spatial rearrangements of chromatin regions and re-organisation of chromatin architecture signifying functional implications as the cell progresses towards a committed cell stage during B-cell development. We first qualitatively investigated if there were any changes in chromatin organisation by doing the comparative analysis of the two simulated structures representing the two different stages of B-cell differentiation. We first plotted the number of compartments in both the cell stages as identified from their respective simulated structural models (Figure 6a).

**Figure 6:**
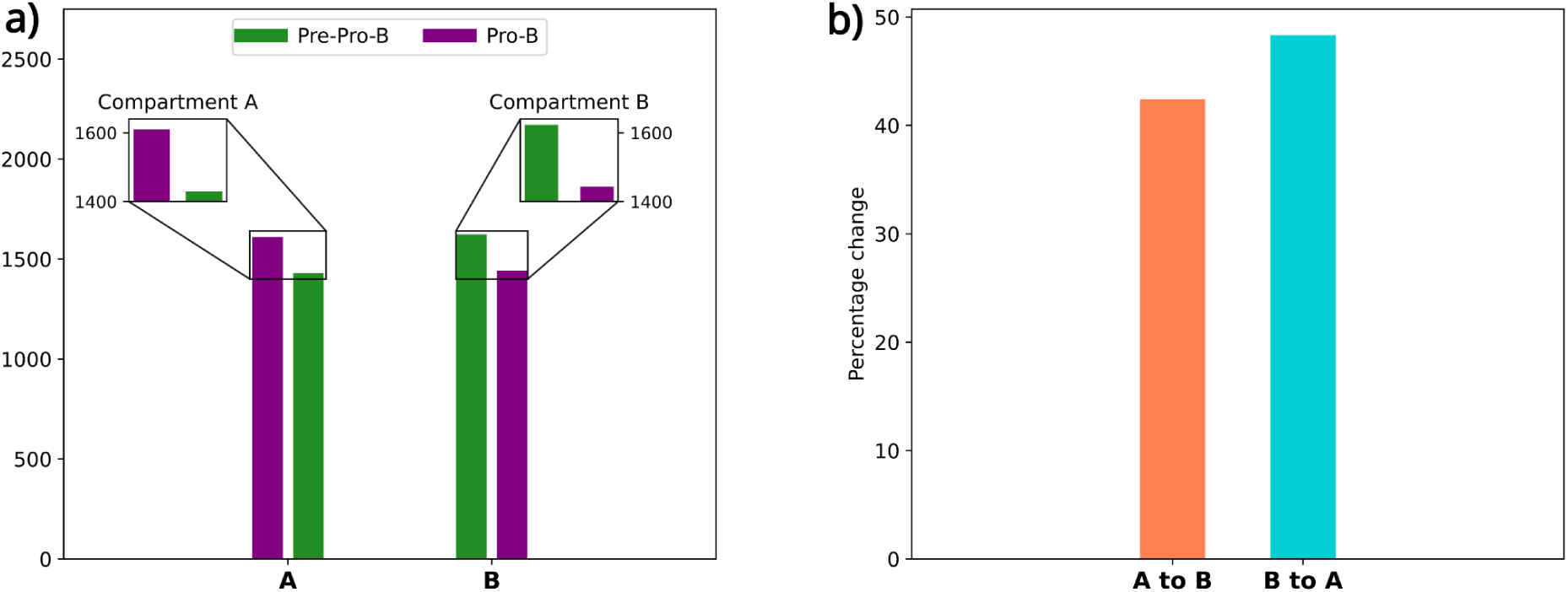
Comparative analysis of simulated structures. (a) Comparison of differential number of compartments in Pre-Pro-B and Pro-B. (b) Compartmental rearrangement from A to B and vice versa during cell differentiation as the cell progresses from Pre-Pro-B to Pro-B stage.

It is observed in Figure 6a that there is indeed a rearrangement of chromatin architecture in Pro-B cells as evident from the difference in the number of A and B compartments between the two simulated structures. This indicates differential transcriptional states of chromatin regions in the two cell types highlighting their contribution towards maintaining the cell identity and also being responsible for governing cellular transitions. To further support it, we quantitatively investigated the number of compartments switching from A to B and B to A compartments and plotted the result in Figure 6b. The result confirms that although small, the chromatin undergoes compartmental switching as the cell differentiates from Pre-Pro-B to Pro-B stage during the B-cell development. We anticipate that this developmental change leads to the activation and repression events of lineage specific and multi-lineage genes, respectively, leading to switching of compartments between permissive and repressive states as the cell transitions towards a committed and differentiated cell stage (i.e. Pro-B) from an undifferentiated stage (i.e. Pre-Pro-B). The small-scale difference is justified because, firstly, this transition from an undifferentiated to a differentiated cell stage is a collective outcome of the differential changes contributed by all the chromosomes of the cell which weren’t considered in our model. Hence, our results show only the contribution of the chromosome under consideration for this study, which is the sub-set of the concerted dynamics brought by the entire genome. Secondly, the two cell stages under consideration are otherwise very similar in their expressions except for the small yet crucial lineage dependent differential expressions. Therefore, the set of differential genes here undergoing the transitions could be very small as compared to the set of other genes maintaining the similar state of expression in both the cell stages; but to be able to detect these changes has proved to be a phenomenal achievement by our model. Within the scope of this study, the model’s performance is highly remarkable as it succeeds in detecting those crucial consequential changes (with limited initial parameters) that were very difficult to detect otherwise. Therefore, we were further interested to investigate those specific regions which underwent the shift in their chromatin states. To do so, we compared the compartmental status of the entire chromatin of both the simulated structures and identified those regions that showed compartmental switching which consequently, contributed to the differential functional state of the cell. In order to quantitatively identify these switched regions, we compared the PC1 values of both Pre-Pro-B and Pro-B cells (top two panels in Figure 7) and identified genomic regions that showed opposite signs in their corresponding PC1 values in the two cell types. The regions shown as green bars in the bottom-most panel in Figure 7 are the regions that switched from either permissive to repressive or repressive to permissive compartments in Pro-B cells. In total, >4% of the regions showed compartmental switching from permissive to repressive (A to B) compartments while >5% of the regions showed the reverse trend in Pro-B cell stage (Figure 6b). These results substantiate that the compartmental switching between A/B compartments correlates to cell type specific genetic switch. To further verify, it would be interesting to know the expression of genes harboured by these switched regions in order to establish functional relevance associated with the switching observed.

**Figure 7:**
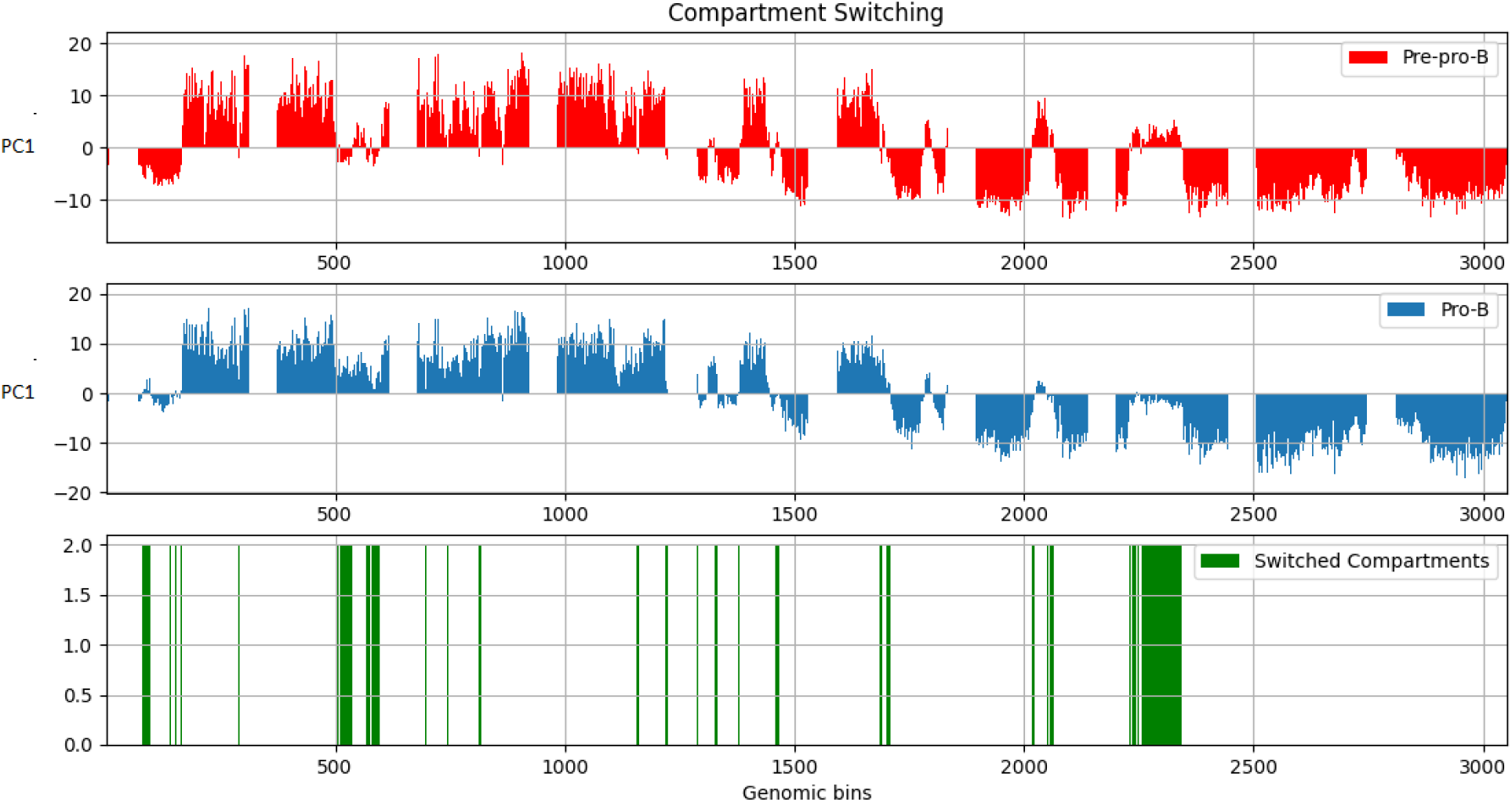
Identification of switched compartments. Comparison of PC1 values of both Pre-Pro-B and Pro-B cells is plotted in the first two panels. The regions shown as green bars in the third panel are the compartments that switched from permissive to repressive and vice versa in Pro-B cells.

### Differential spatial positioning of switched compartments reveals dynamic structural rearrangements in chromatin

We were then interested in investigating changes in the spatial positioning of these switched regions in the two cell types. Within a single chromosomal territory, it has been observed previously that the inner region is comprised of more condensed chromatin domains, while a thin layer of more decondensed chromatin, known as the perichromatin region, can be found around the chromosomal periphery (Fakan and van Driel 2007). Also, the spatial positioning of the genome critically impacts its function and that’s why genomic regions have preferential positioning in 3D space corresponding to their chromatin state. Functionally, the most active genomic regions preferentially lie at the surface of the chromosomal territory while the inactive regions are buried inside, which is also demonstrated in our simulated structures for both cells (Figure 4e and S2e). Since we had observed a shift in the chromatin status of some of the genomic regions of the Pre-Pro-B cell, we were intrigued to investigate the corresponding changes in the spatial positioning in 3D of these switched regions in the two cell types from their respective structural models. With this known phenomenon as the basis of our next analysis, we tried to investigate the distance of these switched regions from the innermost centre, COM of the chromatin. We plotted the mean distance of all the switched bins (both from A to B and B to A) from the COM of the chromatin (Figure 8). It was observed that the distance between the spatial positions of most of the regions switching from permissive A compartment in Pre-Pro-B (blue dots in Figure 8, *top*) to repressive compartment in Pro-B cells (orange dots in Figure 8, *top*) and the centre of mass of the chromatin, reduces in Pro-B indicating a shift in their spatial position towards the interior of the chromosomal territory. Since these regions show switching into repressive compartments, they undergo dynamic spatial rearrangement and move from the periphery towards the chromatin interiors which is also indicative of the preferred position of inactive chromatin state.

**Figure 8:**
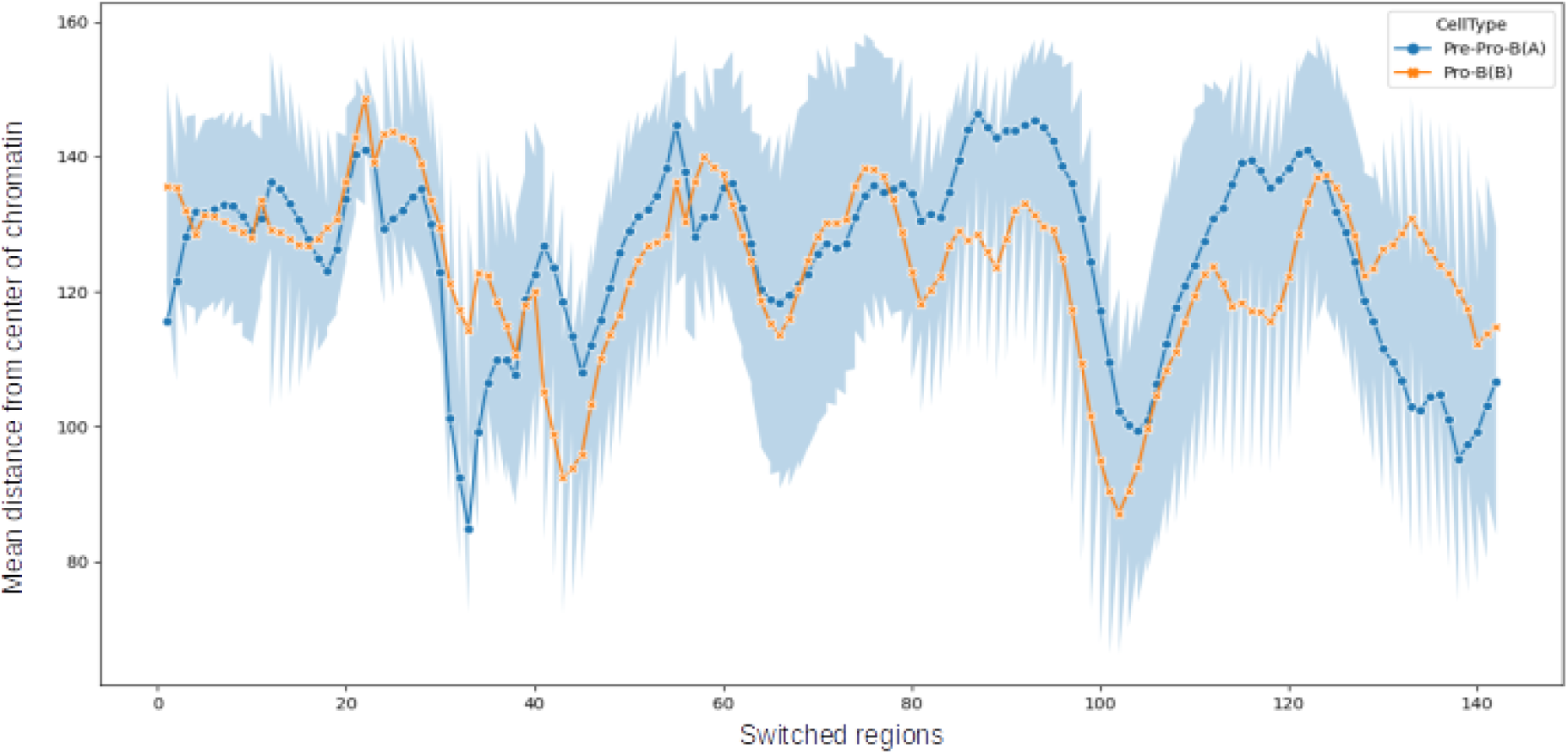
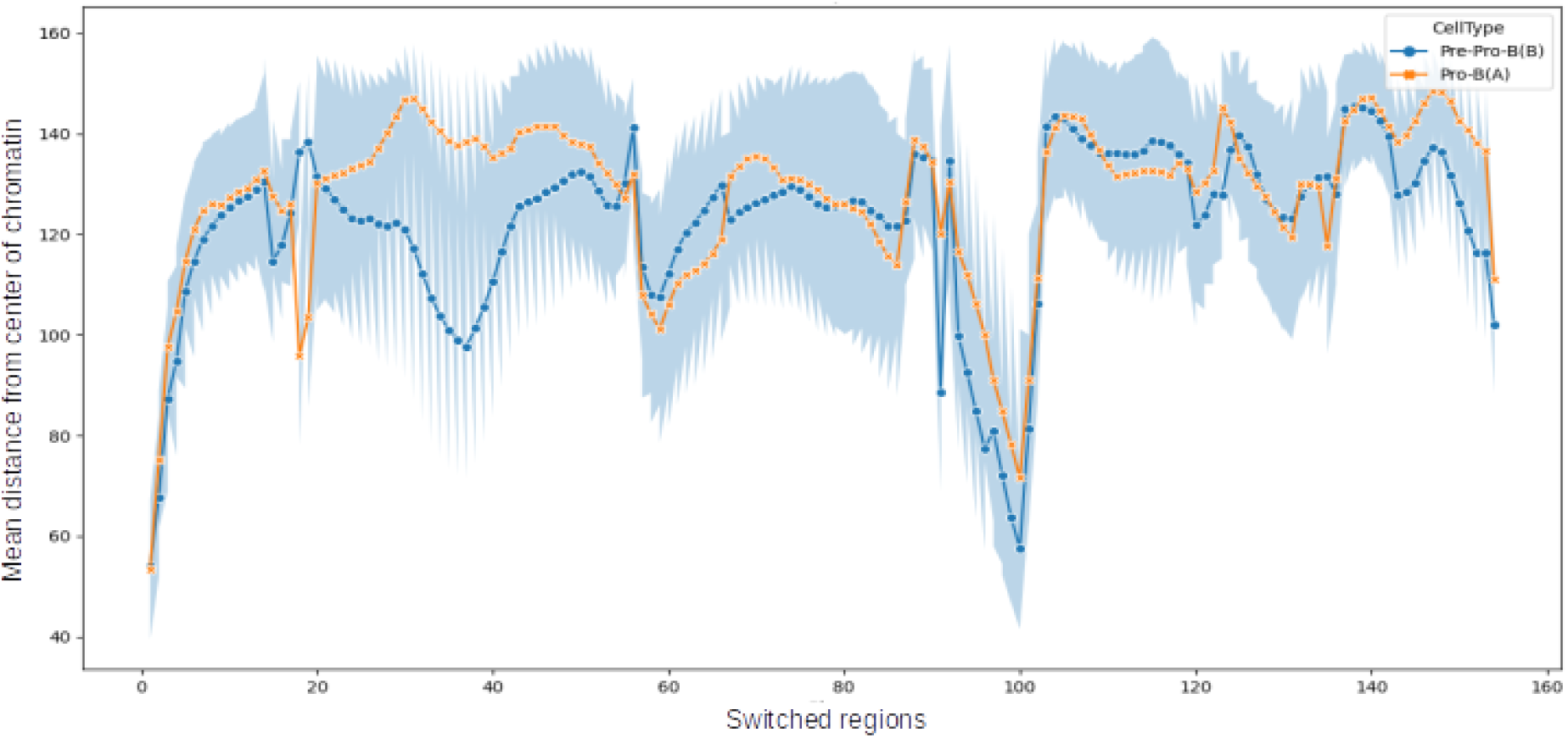
Spatial positioning of regions switching compartments. (a) Compartments switching from A compartment in Pre-Pro-B to B compartment in Pro-B. Blue dots indicate the distance from the COM of regions in the permissive compartment in Pre-Pro-B while orange dots indicate their distance from COM in the switched repressive compartments in Pro-B. (b) Compartments switching from B compartment in Pre-Pro-B to A compartment in Pro-B. Blue dots indicate the distance from the COM of regions in the repressive compartment in Pre-Pro-B while orange dots indicate their distance from COM in the switched permissive compartments in Pro-B.

On the other hand, a reverse trend was observed where an increase in the distances between the regions switching from repressive B compartments in Pre-Pro-B (blue dots in Figure 8, *bottom*) to permissive A compartments Pro-B cells (orange dots in Figure 8, *bottom*) and the COM of the chromatin suggested spatial rearrangement of regions in the repressive compartment residing at the interior, to the permissive compartment shifting towards the periphery of the chromosomal territory which is indicative of the preferential positioning of activated chromatin state. Together, the overall investigation undoubtedly indicates that the genomic regions spatially rearrange themselves depending on their acquired active or inactive status and dynamically move towards their preferential positions within the chromosomal territory. This clearly implies that the chromatin undergoes dynamic structural alterations in the Pro-B cell stage, orchestrating corresponding functional implications resulting in a committed cell stage.

### Degree of compactness of switched regions corresponds to lineage-dependent alterations in chromatin structural framework

Further, in support of our previous findings, we moved forward to investigate if there exists any change in the compactness and folding of the switched regions. The compactness of a region measures the degree of openness or closeness in 3D space which is also associated with the chromatin state and function. In order to examine these, we computed the radius of gyration, R_g_ (a popular metric in polymer-physics) which measures the compactness of a region that also correlates to the accessibility of that region; lower R_g_ value represent a more condensed or compacted state indicating an inactive repressed region while active permissive regions are less compacted and decondensed having a larger R_g_ value providing an easy access to the transcriptional machinery. As a first step to calculate R_g_, we identified and selected regions having a continuous stretch of more than four beads showing compartmental switching. Then we computed the Rg of these regions in both Pre-Pro-B and Pro-B simulated structures. From Figure 9 (*top*), we find that the R_g_ value of most regions in the permissive compartment in Pre-Pro-B structure show a slight reduction when they switch to repressive compartment in Pro-B cell. It is to be noted that the effect is more pronounced and easily visible in regions that are longer in length (regions from 2235 to 2244 and from 2265 to 2343 bead in Figure 9 (*top*)) than the regions of smaller length comprising of 4 beads (regions from 1464 to 146 and from 2060 to 2064), as the measurement of compactness makes more sense as the length of the region Increases. The same holds true in the case of Figure 9 (*bottom*) where an increase in the R_g_ value of most regions switching from repressive compartment in Pre-Pro-B cell to permissive compartment in Pro-B cell was observed. These results clearly show that the compartmental switching is favoured by relative change in the compactness of those regions where active regions in the Pre-Pro-B stage acquire a more compacted structure when they switch into inactive compartments in the Pro-B stage while the inactive regions in Pre-Pro-B open up and attain a comparatively less compacted decondensed structure when they switch to active compartments in Pro-B cells. Hence, we demonstrate a shift in the chromatin structural framework that governs the functional state as the cell differentiates into lineage-specific developmental stages.

**Figure 9:**
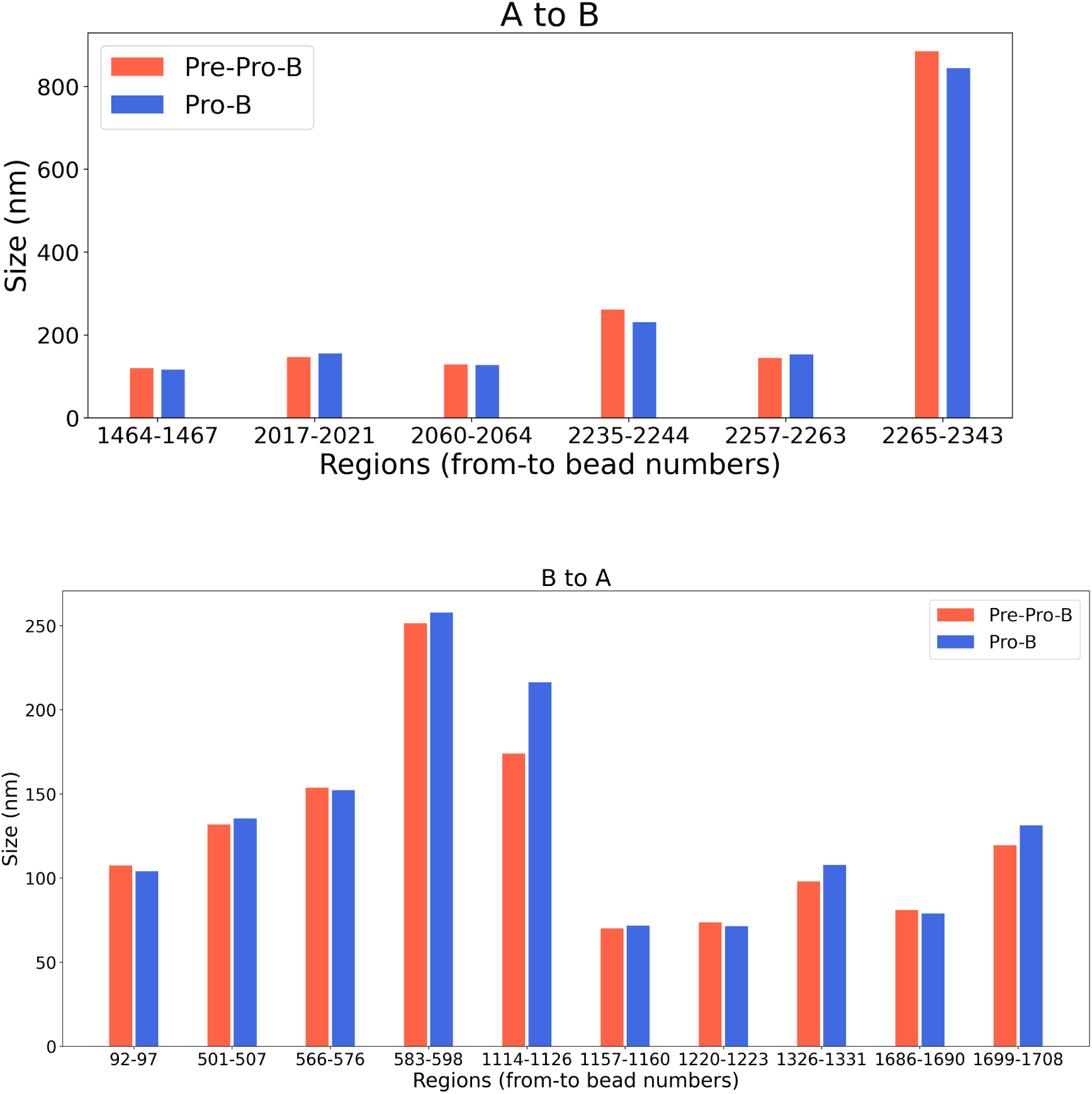
Analysis of compactness of regions that show compartmental switching. (a) Radius of gyration (R_g_) of regions switching from permissive A compartment in Pre-Pro-B to repressive B compartment in Pro-B. (b) Radius of gyration (R_g_) of regions switching from repressive B compartment in Pre-Pro-B to permissive A compartment in Pro-B.

We specifically show the above changes occurring in Ebf1, the master regulator and crucial factor for B-cell commitment (Medina *et al*. 2004, Pongubala *et al*. 2008) through our simulated structures in Figure 10 (*top*). It is evident that the Ebf1 region (red beads) in Figure 10 (*top left*) is a compact region buried in the interior of the chromatin that rearranges itself towards the chromosomal surface and acquires an open chromatin state as it switches to permissive compartment in the Pro-B structure shown in Figure 10 (*top right*). This provides evidence of the structural change in the chromatin organisation framework during differentiation having consequential functional implication of activation of lineage dependent gene, Ebf1, in Pro-B cells confirming B-cell fate commitment. Thus, through our model, we were able to show activation of lineage-dependent genes is related to 3D changes in the structure and architecture of chromatin.

**Figure 10:**
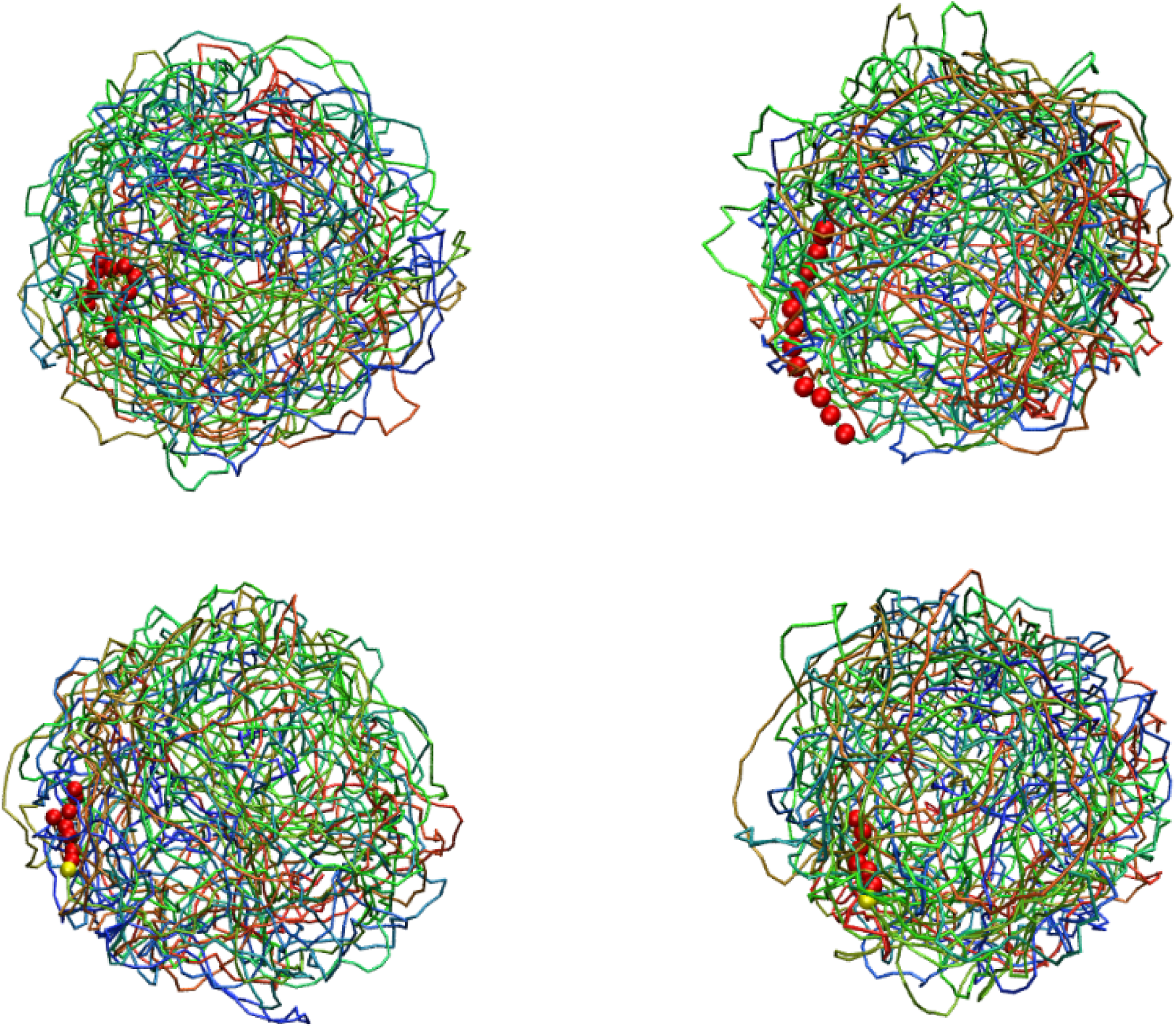
Examples of differential 3D positioning of lineage dependent and alternate-lineage dependent genes. (*top*) 3D position of lineage-dependent gene Ebf1 (in red beads) in Pre-Pro-B (*left*) and Pro-B (*right*) simulated structures. (*bottom*) 3D position of alternate-dependent gene Ccl11 (in red beads) in Pre-Pro-B (*left*) and Pro-B (*right*) simulated structures.

Similarly, Ccl11 is a chemokine gene from the CC subfamily that displays chemo-tactic activity for eosinophils only. It is an eosinophil-specific chemokine that has no significant functional relevance in the B-cell development and hence, is an alternate lineage-dependent gene. From the 3D positioning of Ccl11 (in read beads) in Figure 10 (*bottom)*, it is distinctly evident that 3D spatial positioning of Ccl11 gene shifts from the exterior of the chromatin structure (in the undifferentiated Pre-Pro-B cell stage in Figure 10 (*bottom left*) and buries towards the interior in the committed Pro-B cell stage Figure 10 (*bottom right*). Although small, a relative increase in the compactness of the gene is observed in the simulated structure of Pro-B stage indicating an inactivation event of the function of the gene. Together from these results of both the genes, we confirm the lineage-dependent dynamic structural alterations in the chromatin architecture during B-cell development.

### Prediction & quantitative validation of novel differential genes and their role in maintaining cell identity

So far through our model, we were able to identify and compare the chromatin architectural changes between two cell types during differentiation. From the convincing performance of our model to accurately capture these intrinsic and differential features of the chromatin organisation, we extended its capabilities to predict novel differential regions that weren’t captured in the experiments but showed evident changes in our simulated structures of the two cell types. We further supported our model’s prediction through experimental validations in order to establish this predictive behaviour to our model’s existing features.

We first annotated the genomic regions which showed compartmental switching, with genes from the publicly available data in UCSC (http://genome.ucsc.edu) and other published resources (Smith *et al* 2019) in order to cross-examine their functional roles that can be associated to the observed compartmental switching. All the genes switching compartments from Pre-Pro-B to Pro-B, identified through our simulated structural model of Pre-Pro-B and Pro-B cells are listed in Table S1. The goal was to identify if the predicted switched regions possessed any lineage-specific or alternate-lineage genes that underwent gene activation and gene repression events. Next, we compared this list with the publicly available RNA-seq expression data of both cell types (Heydarian 2014) and identified genes in our results which showed differential patterns in the two cell types that were not captured in the RNA-seq expression data. We call the novel list of these genes as predicted exclusively from the simulated structures (genes in bold in Table S1). Next, we functionally annotated these genes and selected a few genes (Table 1) to be analysed quantitatively through RT-PCR for an experimental validation. We found that the majority of the predicted genes in regions switching from permissive to repressive compartments were related in the developmental expression in alternate-lineage immune cells. For example, genes such as Ccl7, Ccl11 and Ccl12 are not expressed in Pro-B cells but in neutrophils, mast cells, macrophages and other cells of alternate lineages. On the other hand, genes in regions switching from repressive to permissive compartments had characteristic roles in the development and maintaining the identity of B cell and begin to express in the Pro-B cell stage. The same trend has been observed through the results of RT-PCR analysis showing the reduced expression of genes involved in the development of alternate lineages as they switch from permissive to repressive compartments in Pro-B cells (Figure 11 *left*) while B-cell related genes harboured by the regions switching from repressive to permissive compartments are upregulated in Pro-B cell stage (Figure 11 *right*). Through these results, the prediction of gene switching was duly validated and we confirm the predictive potential of our model.

**Figure 11:**
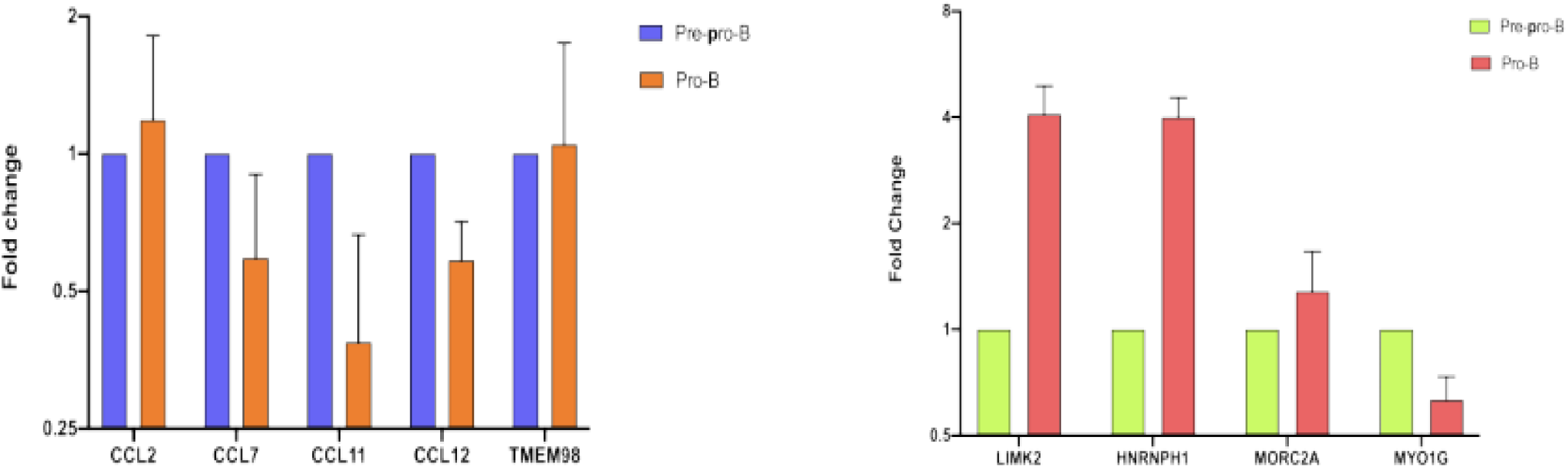
Quantitative analysis of annotated genes by RT-PCR. (***Left)*** Genes predicted to switch from permissive to repressive (A to B) compartments show downregulation in Pro-B cells due to their involvement in the development of alternate-lineages. (***Right)*** Genes predicted to switch from repressive to permissive (B to A) compartments show upregulation in Pro-B cell which is a B-cell committed cell stage and marks the expression of B-cell related genes.

**Table 1:**
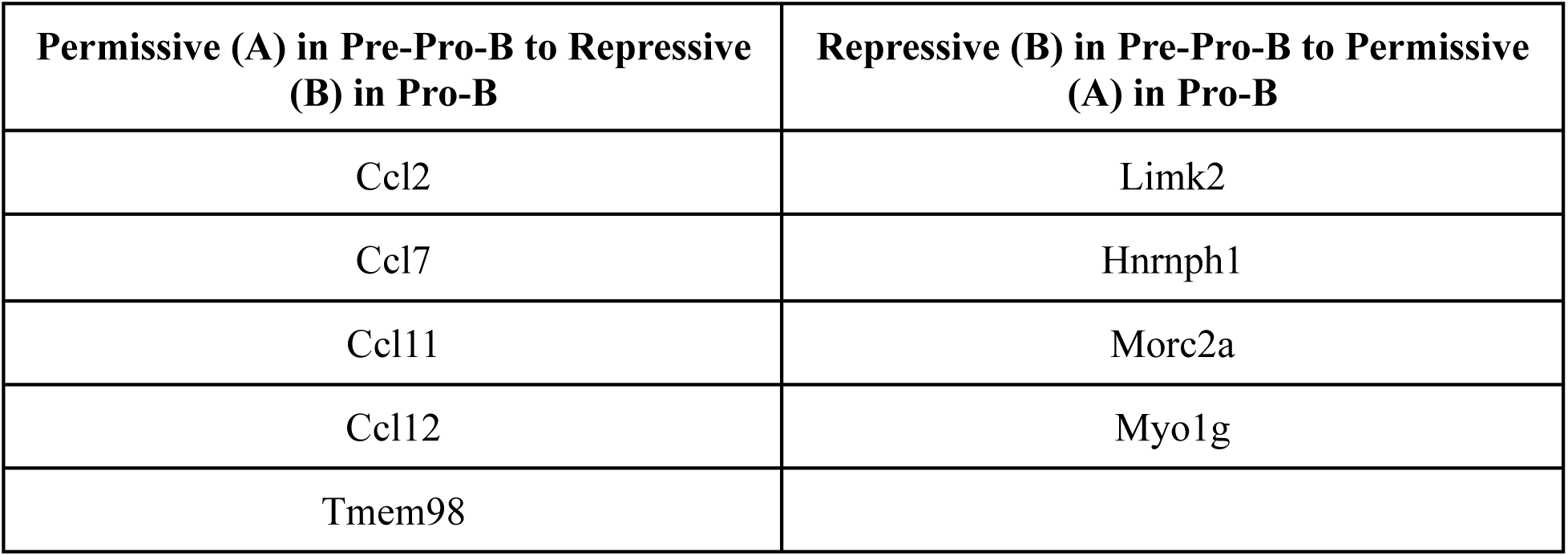
Selected list of genes for experimental validation through RT-PCR.

## Discussions

The complex yet indispensable relationship between chromatin architecture and its impact on the functional state of the cell has been an interest of research for many years in gaining insightful learnings on the underlying mechanisms associated with different cellular functions ranging from cell development, differentiation, maintenance, cell repair etc. This can be better understood by appreciating the three-dimensional organisation of the genome that mediates genomic interactions in 3D nuclear space to bring out the desired cell-type specific functional implications. Hence, a lot of work including experimental and computational studies has been carried out and continues to improve our knowledge on this aspect.

The investigation of these cell-type specific patterns in the context of 3D during the process of cell differentiation was the overall aim of this study. In particular, we looked into the three-dimensional structural architecture and dynamics of chromatin during the formation of B-cells through mechanistic modelling using a combinatorial approach of polymer physics and high-throughput chromosomal conformation capture Hi-C data. We presented a computational model that not only captured the hierarchical structural organisation but also provided mechanistic insights into the spatial rearrangements of chromatin during developing lymphoid lineage cells. Through this study, we were able to show the *spatial dynamics* and 3D transitional rearrangements in chromatin organisation upon cell differentiation, which were not possible through data-driven reconstruction-based modelling approaches that are limited to reconstruction of only static chromatin structures based on the input provided. Also, benetting from our polymer-based predictive approach, we went ahead to make significant differential structural predictions through our simulated structures, which could not be captured in other high-throughput experiments but were detected by our simulated structures.

We show that our simulated structures succeed in independently recapitulating the salient features of different levels of chromatin architecture and additionally, help in identifying the cell-type specific 3D organisation of the chromosome between the two different cell stages considered in our study. In particular, it faithfully reproduced all the considered Hi-C interactions while showing remarkable agreement to chromatin interactions that were not included while generating the initial model structures. Further, we were able to show the intrinsic features of chromatin organisation including folding and local packing as a fractal globule, compartmentalization into permissive A and repressive B compartments and formation of TADs even at the sub-chromosomal scale. The model’s predictions were in agreement for long-range interactions with some amount of noise observed for short-ranged chromatin interactions. These results established the integrity of our model with minimalistic inputs without relying on the proximity-based experimental data. Additionally, in 3D space, we demonstrated through our simulated structures the spatial dynamics and positioning of chromatin into phase separated regions based on their similar chromatin states. Through the mean distances of different regions from the centre of the chromosome, we confirmed that the preferential position in 3D of permissive regions is at the periphery of the chromosomal territory, while repressive regions tend to reside at the chromosomal interiors.

After the successful predictions of the 3D chromatin structures & organisation, we further extended its use in investigating the cell type specific differential changes by performing a comprehensive comparative analysis of the two cell types of the B-cell developmental stages. Our model revealed that chromatin undergoes compartmental switching and dynamic 3D spatial rearrangements during cell differentiation towards B-cell commitment. Although the transitions of lineage specific genes were observed to be small as compared to other genes maintaining the similar state of expression in both the cell stages, yet being able to detect these changes with the help of our model, has proved to be a phenomenal achievement. Within the scope of this study, the model’s performance is highly remarkable as it succeeded in detecting those crucial consequential changes (with limited initial parameters) that were otherwise very difficult.

From the investigations of compactness of switched regions, we showed that the genomic regions acquire an open or closed state depending on their switched active or inactive status and dynamically move in 3D space towards their preferential positions within the chromosomal territory. Our results clearly implied that chromatin undergoes dynamic structural alterations in the Pro-B cell stage, orchestrating functional implications resulting in a committed B-cell stage.

A major advantage of our model is that it further allowed us to make important structural and functional predictions about chromatin rearrangements & folding and its relationship with gene regulation which would not have been detected by simple qualitative examination of the Hi-C data. We were able to predict switching of novel regions from permissive to repressive and vice versa during cell differentiation through our simulated structures. These cell type specific chromatin organisation predictions were further quantitatively validated in vitro. The role of the genes in the predicted regions showing downregulation in Pro-B cells is largely associated with alternate lineage development related events, confirming the cell’s commitment towards B-cell fate, thereby also confirming the reliability and predictivity of our model. This predictive model, thus, presents a significant leap forward in understanding the 3D chromatin architecture and *in silico* study of the 3D chromatin architecture and dynamics of differentiated versus undifferentiated cells during development of lymphoid-lineage cells.

