## Supplementary Information for "Discovery of dynamic changes in 3D chromatin architecture through polymer physics model"

### 1. Figures

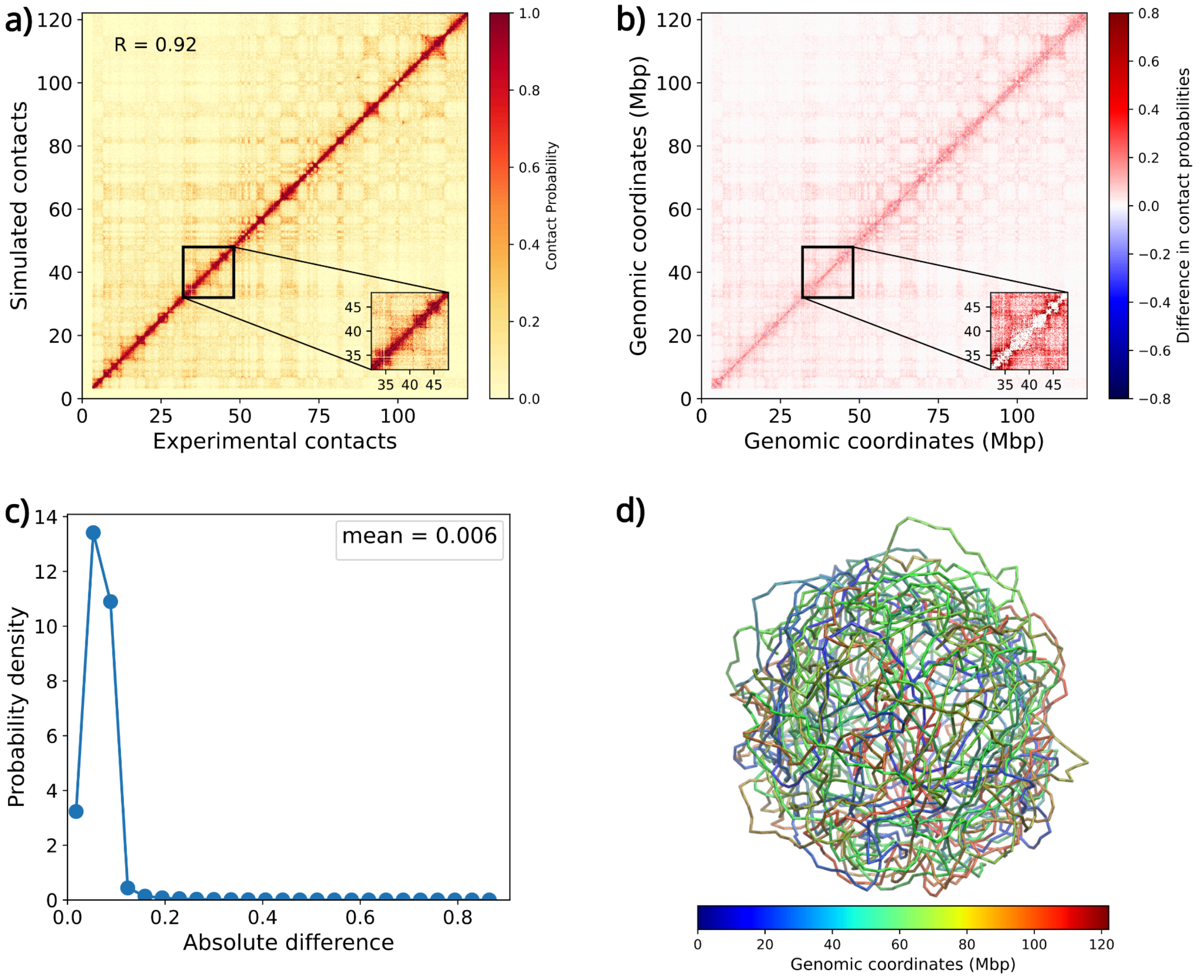


**Figure S1: a)** Comparison of experimental versus simulation-derived probabilities: The heatmap shows the comparison between experimental and simulated contact probabilities maps of chromosome 11 at 40kb resolution. **b)** Difference plot: The heatmap shows the difference between experimental and simulated contact probabilities maps of chromosome 11 at 40kb resolution. The red and blue colours in the colour bar indicate higher contact probability in the experimental data and simulations respectively. **c)** Absolute difference plot: The absolute difference plot shows maximum difference value of 0.006 between experimental and simulation-derived probabilities. **d)** Representative snapshot of a chromosome in the Pro-B cell. The regions of the chromosome have been coloured with respect to their genomic location.

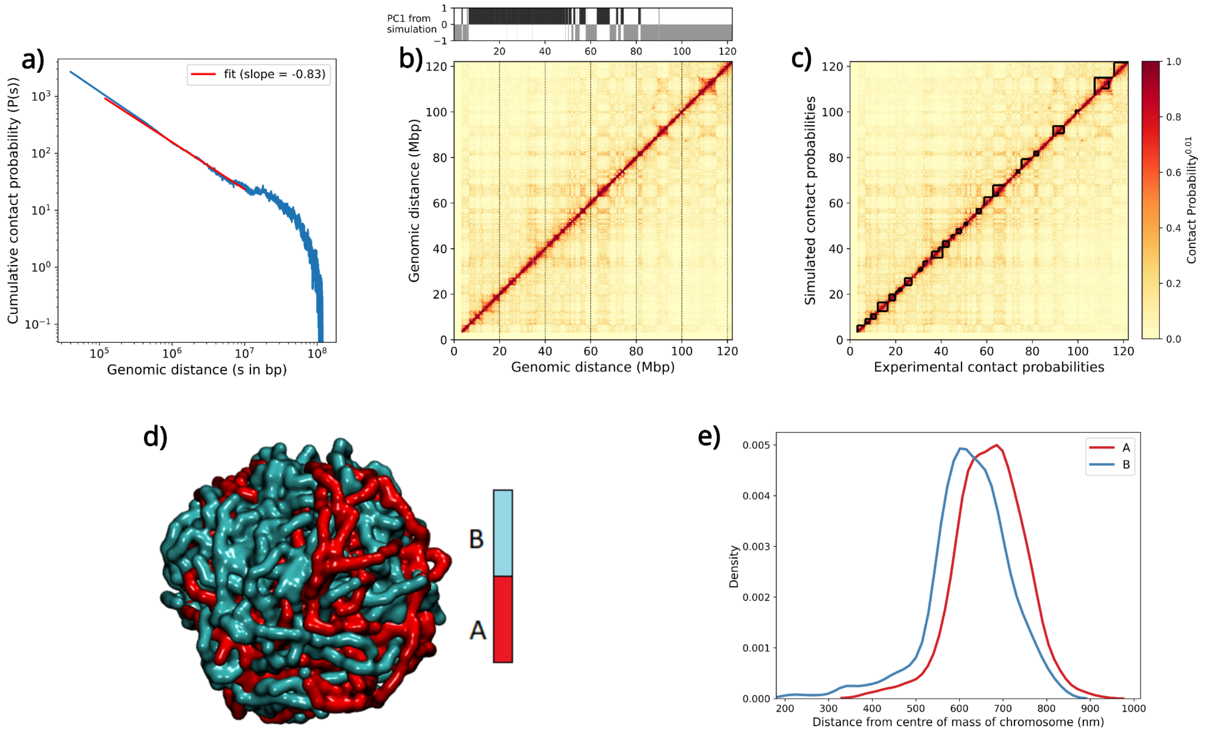


**Figure S2: a) Chromatin folding prediction** Plot of contact probability as a function of genomic distance with a slope (fit shown in red) of -0.83 for Pro-B which is close to the slope of -1.0 for a fractal globule structure. **b) Prediction of chromatin state** Prediction of chromatin states from PCA (PC1 values) of the simulation derived contact probabilities is compared with the heatmap of experimental contact probabilities for Pro-B. The plot shows that the prediction of A compartments (or permissive regions) from the positive PC1 values (black region) in the simulation corresponds to the high contact probabilities in the experimental matrix while the prediction of B compartments (or repressive regions) from the negative PC1 values (grey region) in the simulation corresponds to the low contact probabilities in the experimental matrix. **c) Prediction of Topologically Associated Domains** We have used Armatus (Filippova *et al* 2015) to predict TADs from the simulated (blue triangles) structures and the experimental (green triangles) contacts obtained from Hi-C. The results are shown for Pro-B simulated structures. In this figure TADs which are at least 1 Mbp long have been shown. **d) Phase separation** Phase separation of A and B compartments in the simulated structures for Pro-B cell stage. Similar compartment regions (A compartment in red and B compartment in cyan colour) co-localize leading to a phase separation of active (permissive) from inactive (repressive) regions. **e) Preferential spatial positioning of compartments** Mean distances of A and B compartments from the centre of mass (COM) of the simulated structures of Pro-B. A compartments have larger mean distance from the COM of the chromatin indicating their preferential positioning farther from the centre, at the chromosomal periphery while smaller mean distances of B compartments indicates that their preferential positioning is in the interior of the chromatin, closer to the centre of the chromatin polymer.

### 2. Tables

**Table S1:** List of genes switching compartments from Pre-Pro-B to Pro-B, identified through the simulated structures of the two cell types. The genes in bold are novel predictions of genes exhibiting compartmental switching in Pro-B, exclusively identified in the simulated structural models.

| Permissive (A) in Pre-Pro-B to Repressive (B) in Pro-B | |
|---|---|
| **2210407C18Rik** | 0610010F05Rik, Gm12167, **Myo1g** |
| **4930405D11Rik** | 1700030C12Rik, Gm12184, Nacad |
| **4930507D10Rik** | 1700061J23Rik, Gm12185, **Ntn1** |
| 4930527B05Rik | 1700093K21Rik, Gm12188, **Nudcd3** |
| 5530401A14Rik | 2610024D14Rik, Gm12192, Olfr1393 |
| **Ankfn1** | **4921536K21Rik**, Gm12193, Olfr1396 |
| Asic2 | 4930512M02Rik, **Gm12194**, Olfr56 |
| **Car10** | **8430429K09Rik**, **Gm12195**, **Osbp2** |
| **Ccl11** | **9130017K11Rik**, **Gm12196**, Papolg |
| **Ccl12** | 9130230N09Rik, **Gm12208**, Peli1 |
| **Ccl2** | **9230020A06Rik**, Gm12209, Pex13 |
| **Ccl7** | **9530068E07Rik** Gm12210, **Phykpl** |
| **Ccl8** | **9930111J21Rik1**, **Gm12235**, **Pik3ip1** |
| **Cox11** | Actr2, **Gm12301**, **Pla2g3** |
| **Fam183b** | Aftph, **Gm12303**, **Psme2b** |
| **Gm11207** | Ahsa2, **Gm12304**, Pus10 |
| Gm11416 | **Atox1**, **Gm12305**, Rab1a |
| Gm11417 | B3gnt2, **Gm12592**, Rack1 |
| **Gm11419** | **C78197**, Gm16170, Rasgef1c |
| **Gm11494** | **Canx**, **Gm16518**, Rel |
| **Gm11498** | **Cby3**, **Gm20169**, Rnf130 |
| **Gm11500** | Ccm2, Gm20456, **Rnf185** |
| **Gm11501** | Cct4, Gm22600, Rpl12-ps2 |

| | |
|---|---|
| **Gm11502** | Cct4, Gm22753, Rufy1 |
| **Gm11506** | Cep68, Gm22807, Selenok-ps1 |
| **Gm11511** | **Cfap52**, Gm22990, **Selenom** |
| **Gm11512** | Cnot6, Gm23114, Sertad2 |
| **Gm11516** | Col23a1, Gm23492, Slc1a4 |
| **Gm12251** | Commd1, Gm23582, **Slc35e4** |
| **Gm12252** | Cyp2, Gm23681, **Slc36a1** |
| **Gm12253** | **Dusp18**, Gm23772, **Slc36a1os** |
| **Gm12254** | Ebf1, Gm23813, **Slc36a2** |
| **Gm12255** | Efcab9, Gm23827, **Slc36a3** |
| **Gm12570** | Ehbp1, **Gm24013**, **Slc36a3os** |
| **Gm17268** | **Eif4enif1**, Gm24313, **Smtn** |
| **Gm22599** | Eml6, Gm24398, Snora5c |
| **Gm22702** | Fam161a, **Gm24439**, Snord95 |
| **Gm22762** | Fam71b, Gm24917, Snord96a |
| Gm24612 | Fbxw11, Gm25296, Stk10 |
| Gm24856 | Fstl4, **Gm26157**, **Stx8** |
| Gm25113 | **G3bp1**, Gm26253, **Tbc1d9b** |
| Gm31522 | Gas7, **Gm26393**, Tbrg4 |
| Hlf | Gfpt2, **Gm27194**, **Tcn2** |
| Kif2b | **Gm10428**, **Gm27517**, **Trim41** |
| Lypd8 | Gm11186, **Gm27624**, Trim7 |
| **Lypd8l** | **Gm11189**, **Gm27640**, **Tug1** |
| **Lypd9** | **Gm11944**, **Gm27937**, Ugp2 |
| **Myo1d** | **Gm11945**, Gm28048, Usp34 |
| **Olfr224** | **Gm11948**, Gm30942, **Usp43** |
| **Olf30** | **Gm11949**, **Gm33351**, Vps54 |
| **Olfr311** | **Gm11950**, Gm3718, Wap |
| **Olfr312** | **Gm11951**, Gm40824, Wdpcp |

| | |
|---|---|
| **Olfr313** | **Gm11952**, Gm47279, **Wsb2-ps** |
| **Olfr314** | **Gm11973**, **Gm51877**, Xpo1 |
| **Olfr315** | Gm11998, Gm5431, **Zfp287** |
| **Olfr318** | Gm12030, **Hnrnph1**, Zrsr1 |
| **Olfr319** | Gm12031, **Hspa4** |
| **Olfr320** | Gm12034, I47 |
| **Olfr322** | Gm12035, **Inpp5j** |
| **Olfr323** | Gm12036, Irgm1 |
| **Olfr324** | Gm12037, Itk |
| **Olfr325** | Gm12038, Lcp2 |
| **Olfr326-ps1** | Gm12039, Lgalsl |
| **Olfr328** | Gm12040, **Limk2** |
| **Olfr329** | Gm12041, Mapk9 |
| **Olfr329-ps** | Gm12042, Mdh1 |
| **Olfr330** | Gm12043, Med7 |
| **Olfr331** | Gm12044, Mgat1 |
| **Olfr332** | Gm12055, Mir1933 |
| **Olfr333-ps1** | Gm12056, Mir340 |
| **Spaca3** | Gm12057, **Mir3470a** |
| **Stxbp4** | Gm12058, **Mir6406** |
| **Tmem132e** | Gm12061, **Mir804** |
| **Tmem98** | **Gm12062**, **Morc2a** |
| **Tom111** | Gm12158, **Mup-ps22** |
| **Trim58** | |

#### 3. Determining Bead Size and the Radius of Confinement of each Polymer

##### a. Determining Bead Size, σ

To determine the bead size, we assume that the volume of chromosome 11 having L basepairs ($V_L$) *in vivo* is equal to the volume of the modelled chromosome 11 *in silico*.

Volume of the polymer *in silico* is computed as volume of one bead × total number of beads in the polymer. Therefore, if $4/3\pi(\sigma/2)^3$ is the volume of one bead with diameter σ and N is the total number of beads in the polymer, $V_L$ *in silico* can be written as

**Volume of one bead x Number of beads = Volume of chromosome of length L bp**

$$4/3\pi(\sigma/2)^3 \times N = V_L \qquad \text{...(i)}$$

Now, to compute the volume $V_L$ *in vivo*, we first compute the volume of 1 bp. $V_L$ can then be derived as volume of 1bp × L bp with an assumption of uniform volume of each basepair. Since the entire genome of total length *G* bp occupies a volume denoted by $V_{genome}$, we assume that 1 bp will effectively occupy a volume $V_{genome}/G$ and therefore, chromosome with L bp will occupy a volume $(V_{genome}/G) \times L$ i.e. $V_L$ *in vivo* can be written as

$$V_L = (V_{genome}/G) \times L \qquad \text{...(ii)}$$

Here, if 0.1 is the volume fraction where the volume of the genome occupies 10% of the nuclear volume $V_{nucleus}$, then, we can derive $V_{genome}$ as $\mathbf{V_{genome} = 0.1 \times V_{nucleus}}$ and substitute it in equation (ii) as

$$V_L = (0.1 \times V_{nucleus}/G) \times L \qquad \text{...(iii)}$$

where $V_{nucleus}=4/3\pi(d_{nucleus}/2)^3$ with the nuclear diameter, $d_{nucleus}$ = ~7μm for lymphocytes since in normal situations, the coarse, dense nucleus of a lymphocyte is approximately about 7μm in diameter (Abbas AK, Lichtman AH (2003). *Cellular and Molecular Immunology* (5th ed.). Saunders, Philadelphia). Substituting $V_{nucleus}$ in equation (iii), we obtain $V_L$ *in vivo* as

$$V_L = (0.1 \times 4/3\pi(d_{nucleus}/2)^3/G) \times L \qquad \text{...(iv)}$$

Equating the $V_L$ *in vivo* from equation (iv) and $V_L$ *in silico* from equation (i), we get

$$4/3\pi(\sigma/2)^3 \times N = (0.1 \times 4/3\ \pi\ (d_{nucleus}/2)^3/G) \times L$$

Simplifying that gives us

$$\boxed{\sigma = d_{nucleus}\ (0.1 \times L / G \times N)^{1/3}}$$

For $d_{nucleus}$ = 7μm,

L= 122082543 bp,

G = 2 x haploid = 2 x 2725521370 bp and

N = L/40kbp = 3053 beads, we obtain

**σ = 63.13 nm**

### b. Determining Radius of Confinement, $r_{conf}$

Based on the fact that the total genome has a volume fraction of 0.1 within the entire nuclear volume enveloping that genome, i.e $V_{genome}$ = 0.1 × $V_{nucleus}$; we can assume that $V_L$ would also have a volume fraction of 0.1 within its chromosomal territory defined as $V_{confinement\_for\_L}$. That is to say,

$$\mathbf{V_L = 0.1 \times V_{confinement\_for\_L}}, \quad \ldots(v)$$

where $\mathbf{V_{confinement\_for\_L} = 4/3\ \pi\ (d_{confinement\_for\_L}/2)^3}$

Substituting $\mathbf{V_{confinement\_for\_L}}$ in (v), and then equating the resulting $V_L$ *in silico* to $V_L$ *in vivo* in equation (iv), we obtain

$$\mathbf{0.1 \times 4/3\ \pi\ (d_{confinement\_for\_L}/2)^3 = (0.1 \times 4/3\ \pi\ (d_{nucleus}/2)^3/G) \times L}$$

Simplifying it, we get

$$\mathbf{(d_{confinement\_for\_L})^3 = (d_{nucleus}{}^3/G) \times L}$$

$$\mathbf{d_{confinement\_for\_L} = d_{nucleus} \times (L/G)^{1/3}}$$

$$\boxed{\mathbf{r_{confinement\_for\_L} = d_{nucleus}/2 \times (L/G)^{1/3}}}$$

With values of $d_{nucleus}$ and L defined for chromosome 11 and G mentioned above, we calculated $r_{confinement-for-L}$ or $r_{conf}$ = 986.57nm i.e. 986.57nm/63.13nm = 15.6σ. Therefore, $\mathbf{r_{conf}}$ **= 15.6σ**

### 4. Defining the force fields

The force fields were defined by defining the following potentials:

(i) The bonded interaction potential $V_b(r_{ij})$ between consecutive beads *i* & *j* separated by distance $r_{ij}$ was defined as strong harmonic springs given by the equation

$$\mathbf{V_b(r_{ij}) = 1/2\ k_b(r_{ij}-r_0)}$$

with the equilibrium bond length $r_0$ = 1σ and a strong $k_b$ = 300kJ $mol^{-1}$ $σ^{-2}$.

(ii) The angular potential ($U_{angle}$) between three consecutive beads was defined as

$$\mathbf{U_{angle} = K_a\,[1-\cos(\theta-\theta_0)]}$$

where $k_a$ = 2.0 , $\theta_0$= 180 in order to provide rigidity to the polymer and reduce the possibilities of unwanted bending that can give rise to overlaps between beads.

(iii) Since we assume that Hi-C interactions takes care of the attractive interactions, the repulsive non-bonded interactions were defined as the repulsive term of the Lennard-Jones potential, **$V_{LJ}$ (r) = $c^{12}/r^{12}$ − $c^6/r^6$ with $c^6$ = 0.0** and $c^{12}$ = 1.0 kJ mol $^{-1}$. The interactions between all the non-consecutive bead pairs have been permitted purely via this repulsive potential.

### 5. Derivation of simulation time scales

Simulations were run for the total number of timesteps = $2\text{x}10^6$ timesteps where 1 timestep (ts) = 0.002**τ**. The value of **τ** is calculated as **τ** = $3\pi\eta\sigma^3/k_BT$

where η (viscosity of water) = $10^{-3}$Pa sec,

$1k_BT = 4\text{x}10^{-21}$ J and

σ = 63.13nm

which results in **τ** = 0.593ms. Therefore, the total time for which the simulation ran for each of the configuration was: total timesteps x 1 timestep = ($2\text{x}10^6$ ) × (0.002 × 0.593 × $10^{-3}$ secs) = 2.372s
